# Sampling Function-Related Metastable States of Proteins With DASH

**DOI:** 10.64898/2025.12.24.696448

**Authors:** Jinyin Zha, Zhen Zheng, Jie Zhong, Weihua Wang, Qiao Li, Qiancheng Shen, Mingyu Li, Chengwei Wu, Qingjie Xiao, Qiuhan Ren, Nuan Li, Hao Zhang, Xinyi Liu, Wenming Qin, Li Feng, Jian Zhang

## Abstract

Rational discovery of function-specific protein modulators as well as activity-enhanced engineering proteins underscore the need to identify function-related metastable states (FMSs) of proteins. However, current experimental and computational methods struggle to generate these states directly from their native state (NS), likely because the NS → FMS transition is non-spontaneous. To address this challenge, we introduce **D**eep learning guided **A**daptive sampling with seed **S**election and **H**opping (DASH), integrating both deep learning and physical functions to guide molecular dynamics (MD) simulation towards FMS. DASH successfully sampled NS → FMS transitions in 18 cases across two tasks: protein activation and cryptic allosteric site opening. DASH combined with secondary-structure collective variables is further able to sample folding of disordered regions. Compared to existing methods, DASH demonstrates superior performance while requiring comparable simulation time. Crucially, we applied DASH to sample new conformations of proteins, by which revealing new folding states, previously unknown allosteric sites, and potential activators. These predictions have been further verified in wet-lab experiments and crystal structures determinations. Collectively, our framework provides a robust strategy for function-specific pharmacy research and could accelerate future drug discovery efforts.

## Introduction

Proteins are dynamic, switching among a predominantly populated native state (NS) and several less populated metastable states (MSs) ^1–3^. Among these MSs, some could be stabilized to become a dominant conformation via ligand binding or mutation occurrence, during which the energy landscape is reshaped (Fig. 1a). Such shifts might enhance^4^, impair^5^, or even give rise to protein functions^6^. Therefore, structural identification of these function-related MSs (FMSs) is critical for rational discovery of function-specific modulators^7–9^ as well as engineering proteins with improved activity^10–12^. However, experimental characterization of MSs as well as FMSs is considerably harder than that of NS due to their shorter life time, and often requiring serendipitous discovery of stabilizing agents or mutations beforehand^13, 14^. This challenge calls for developing *in silico* methods for identification of MSs and especially FMSs.

**Fig. 1.**
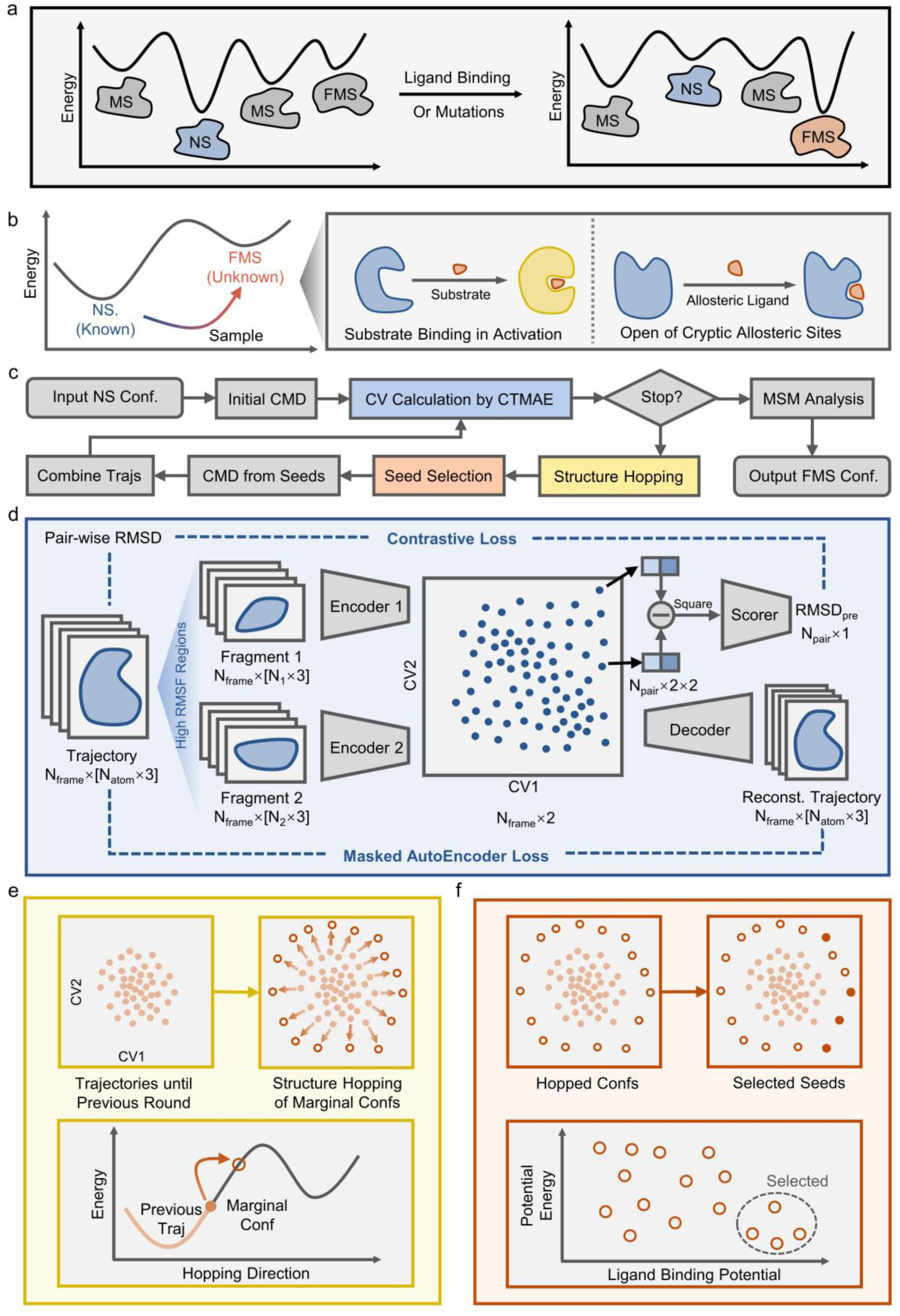
Overview of DASH. **a.** Illustration of native state (NS), metastable state (MS) and function-related metastable state (FMS) of a protein. **b.** Problem description: sampling the function-related metastable state (FMS) conformation from native state (NS) conformation, including protein activation and cryptic allosteric sites opening **c.** Workflow of DASH. **d.** DASH uses a combined contrastive learning and masked autoencoder (CTMAE) framework for CV calculation. **e.** Structure hopping. Marginal conformations are hopped on the CV space to enlarge tendencies of conformational transitions and overcome energy barriers. **f.** Seed selection. Hopped structures are selected based on their potential energy and ligand-binding potential. This step helps filter out structures more likely to approach FMS, in order to realize selective boosting of directions towards FMS.

Deep learning models^15, 16^ have revolutionized protein structure predictions, but recent studies show that they tend to output NS-like conformations rather than MS^17, 18^. Molecular dynamics (MD) simulation is a powerful tool to study protein conformational changes. Ideally, one could start an MD simulation from an easier-fetched NS to generate a variety of MSs, and then to cherry-pick FMS based on trajectory information and prior knowledge. However, the fact is that transitions between distinct conformations often occurs on prohibitively long timescales^19–21^. To tackle this limitation, a series of enhanced sampling methods have been developed^22^. Among these, collective-variable-based (CV-based)^23–25^ and transition-pathway-based^26–28^ techniques are highly efficient as they directly accelerate the simulation on transition-related directions. Nevertheless, these techniques are more suitable for sampling transition pathway between two known conformations, because it is challenging to define sampling directions when destination conformation is unknown. In contrast, methods like Gaussian-accelerated MD (GaMD)^29, 30^, replica-exchange MD (REMD)^31, 32^, and adaptive sampling^33, 34^, which do not require predefined sampling directions, are better suited for exploring conformational space from a single state. Recently, approaches combining CV-based methods and adaptive sampling have emerged^35–41^. These involve initially defining approximate CVs, followed by iterative refinement based on accumulated simulation data. However, these methods were mainly applied to sample NS from FMS. The concern for the reversed application (sampling FMS from NS) lies in their uncontrollable boosting directions. Sampling from the higher-energy FMS towards the lower-energy NS is inherently guided via energy descending^42^. Conversely, starting simulations from the lower-energy NS lacks such guidance, potentially driving the system towards nonsense MSs or even unphysical conformations rather than the target FMS. Besides, since NS → FMS is unspontaneous, transition sampling would face a much higher energy barrier^43^.

To achieve FMS-directed sampling from *only NS known*, we introduce here **D**eep-learning-guided **A**daptive sampling with seed **S**election and **H**opping (DASH). DASH iteratively generates transition-related CVs via deep learning, overcomes energy barrier through structure hopping along these CVs, and aims at FMS by selecting hopped seed structures based on physical functions (Fig. 1d-f). DASH has realized direction-free sampling of protein activation and opening of cryptic allosteric sites. The algorithm was extensively benchmarked on 18 cases and its performance was compared with other non-direction-based enhanced-sampling methods. Besides, DASH could be cooperated with direction-known methods to realize sampling of the folding of intrinsic disordered regions (IDR) (Fig. 1b). Moreover, the algorithms were further applied to predict novel activators, allosteric sites and foldable IDRs. The predictions were subsequently validated by wet-lab experiments.

## Results

### Overview of DASH

The main idea behind DASH is that, although conformational transitions exist in prohibitively long timescale, the *tendency* of transitions could exist in short simulations. Therefore, DASH aims to selectively amplify those tendencies more-likely towards FMS on-the-fly of MD simulations.

In practice, DASH operates through an iterative workflow (Fig. 1c). Initially, a short trajectory is generated by conventional molecular dynamics (CMD). Collective variables (CVs) are then calculated using the contrastive learning and masked autoencoder (CTMAE) model (Fig. 1d). The CV-space represents a collection of tendencies of transitions. After that, conformations are uniformly selected from the margin of the CV-space, and hopped via displacing their CV values away from the center (Fig. 1e). This process would amplify any possible transitional tendencies, so that a physics-based filter is followed to select hopped conformations more likely to approach FMS (Fig. 1f). The selected conformations are used as seeds for a new round of CMD simulations, CV calculations, structure hopping and seed selection. Finally, the iteration is over and the trajectory is analyzed with Markov-state model^44^ (MSM).

In the next two sections, we will discuss and benchmark the three crucial designs in DASH: CTMAE and structure hopping and seed selection.

### CV calculations with CTMAE

In CTMAE (Fig. 1d), protein coordinate sets are processed through N_CV_ independent neural network (NN) encoders to generate N_CV_ CVs, composing a reduced representation. The design of multiple encoders not only might reduce correlations among CVs, but also allows different inputs for N_CV_ CV calculations: either whole protein or spatially-clustered fragments with higher root-means-square fluctuation (RMSF) values. Fragmentation is applied here to further isolate motions in different domains. For simplicity, we set N_CV_=2, although applying more CVs has been found to better describe protein motions^40^.

The subsequent structure hopping requires that CVs should both well preserve transitional tendencies and place transitional-pioneering conformations at the margin. Therefore, we trained the NNs on both masked autoencoder (MAE) and contrastive-learning (CT) tasks. MAE employs a decoder to reconstruct complete protein coordinates, which have been proven well-preserving non-linear protein motions^35, 36, 45, 46^. CT is inspired by SimCLR^47^, taking squared deviations of pairs of CV coordinates as input and employs a weak NN scorer to regress the root-mean-square deviation (RMSD) of corresponding protein frames. CT is designed to flexibly encode structural deviations into CV-space and therefore could place transitional-pioneering conformations at the margin, meeting the requirement of subsequent hopping of marginal structures.

We evaluated motion preservation by micro-/macro-state consistency and secondary-structure-state discrimination, while evaluated the quality of marginal conformations by intra-margin and margin-to-all-frames RMSD. The evaluation is carried on 15 differentiated trajectories and compared with 6 other methods (Data S1), with indicators normalized per trajectory. As shown in Fig. 2a, CTMAE demonstrate well performance in all metrics, whereas other methods excel in fewer areas: PCA^48^ and IsoMap^49^ achieve better marginal quality; autoencoder-based^35^ methods do well in motion preservation and TICA^50^ excel at separating different secondary-structure states. Weight adjustments of CT and MAE suggest that CT contributes more on generating better marginal structures whereas MAE contributes more to retaining motions (Extended Data Fig. 1a). Our NN utilizes a simple MLP encoder, while nowadays graph encoders have gained prominence in protein modeling^51–53^. We replaced our encoder with a variety of graph neural network (GNN), but no significant improvement was found (Extended Data Fig. 1b), possibly because only protein coordinates are used as input and MLP is sufficient to handle them. Instead, a great increase was found in time and memory when using GNNs for seed hopping. Overall, the above results suggest our CTMAE model could offer high-quality CVs for seed hopping.

**Fig. 2.**
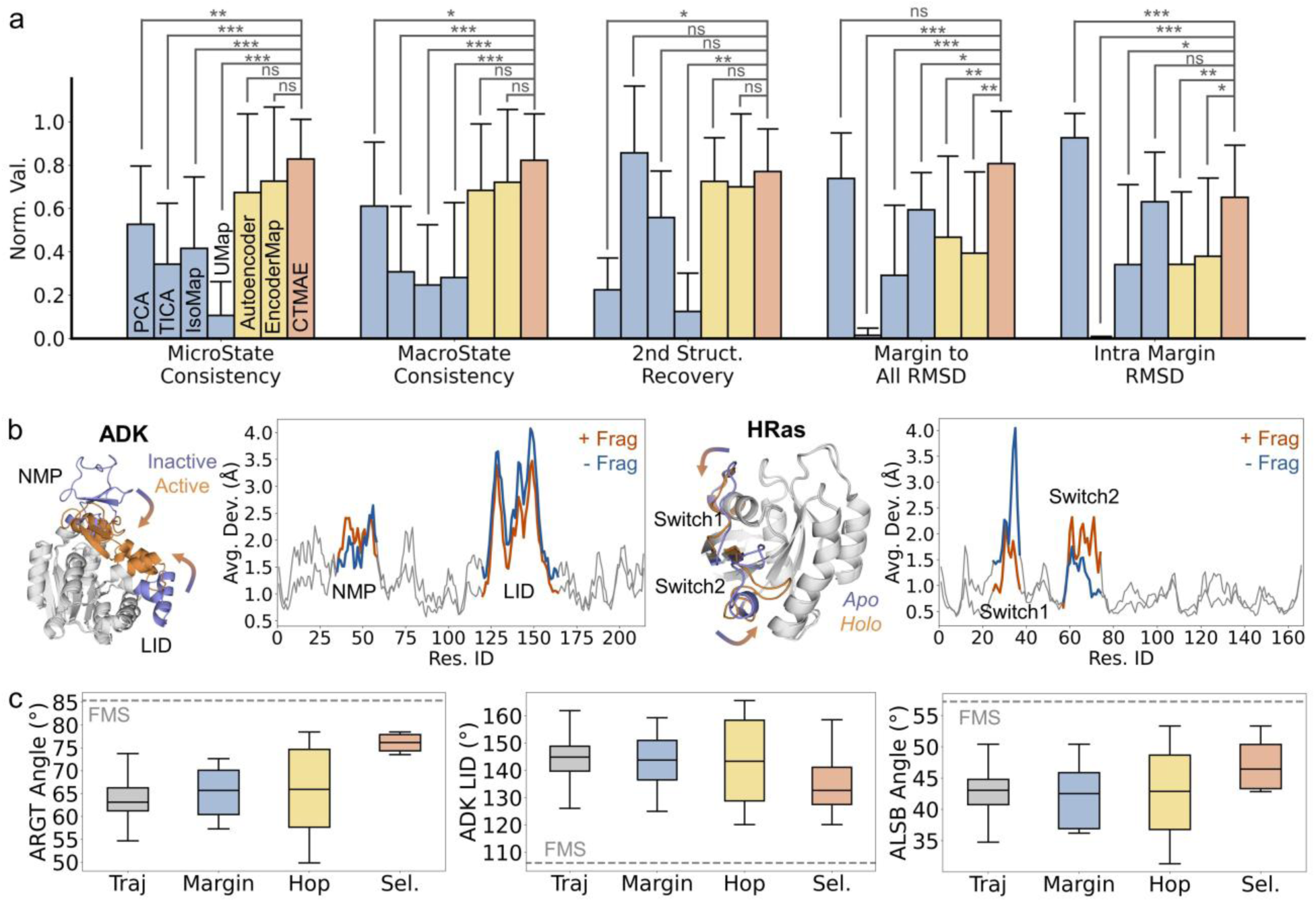
Benchmarking each module in DASH. **a.** Comparison of CTMAE and other dimension reduction algorithms on preservation of thermodynamics (first three indicators) and seed-structure diversity (last two indicators). N=15. Error bars represent standard deviations. Statistics analysis are analyzed using Student’s t-test (normal and homo-variance) or Mann-Whitney test (otherwise). N=10. *: p<0.05. **: p<0.01. ***: p<0.001 **b.** Comparison of multi-domain residues deviations when using / not using the fragmentation strategy in CTMAE. **c.** Effect of seed hopping and seed selection in sampling function-related metastable state (FMS) conformations. 10000 frames in trajectory, 20 margin and hopped structures and 4 selected structures. Box plot represents the range between 25% and 75% quantile, with line on it represents the median value. Whiskers of box plot represents 1.5 times of inter-quantile range.

The rationale for isolating motions of different domains with fragmentation is that, in some NS → FMS transitions, multiple domains move, like ADK^4^ (NMP and LID) and HRAS^54^ (switch 1 and switch 2), so that transitions in different domains could be boosted individually. We generate CVs from 10 ns CMD simulations of NS ADK and HRAS using CTMAE, both with or without the fragment strategy, and perform hopping of marginal structures. As shown in Fig. 2b. The overall movement is not significantly changed after fragmentation. Nevertheless, the deviations of the two conformation-changing domains are more balanced, preventing that only one domain changes significantly while the other changes minimally across sampling iterations.

### Structure hopping and seed selection aids FMS-directed boosting

Structure hopping amplifies transitional tendencies learned by CTMAE (Fig.1e). This is realized by performing step-wise steered molecular dynamics (SMD)^55^ to gradually pull marginal conformations away from the CV space center. This process is designed to overcome high barriers in conformational transitions. Since this process could amplify any transitional tendencies, a subsequent seed selection module (Fig. 1f) is applied to pick out hopped structures that resemble more FMS, which will be used as seed structures for a new round of CMD simulations. In this article, we use potential energy and ligand-binding-potential for seed selection, motivated by the fact that unphysical conformations are energetically high^56^ and ligand-binding-induced^2^ energy shift is considered here. We agree that these filters are not necessary and sufficient conditions of FMS, but at least they could pick out conformations more likely to be closer to FMS.

To evaluate seed hopping & selection, we performed 10 ns simulation of 3 proteins from NS, calculated CVs with CTMAE, and performed seed hopping & selections on marginal conformations in CV space. As illustrated in Fig. 2c, structure hopping effectively broadens the conformational distribution, but such expansion directs some conformations closer to the FMS while moving others further away, proving that structure hopping only provides amplification on any possible tendencies. Importantly, the subsequent selection step filters out seed structures that are closer to the FMS. We have tested other filters including solvent-accessible surface areas and volumes in potential binding regions. (Extended Data Fig. 1c) The performance is worse than the combined potential energy and binding potential filter. The above results indicate that the combined hopping & selection strategy could progress the system towards the desired metastable states.

Having benchmarked the building blocks of DASH, in the following sections, we would introduce three applications of DASH in sampling FMS conformations of proteins.

### Sampling active states from inactive states

During activation, many proteins would undergo significant conformational changes when they bind their substrates^2^. The bound active-state conformation could be viewed as an FMS (Fig.3a). We collected ten such cases and used 30-round DASH to sample the active-state conformation from the inactive state. The first round consisted of a 10-ns conventional molecular dynamics (CMD) simulation, while in each following round, four 2.5-ns CMD simulations were performed from four selected seeds, resulting in a total simulation time of 300 ns per case. An example DASH simulation of glutamine binding protein^57^ (GLNH) is shown in Fig.3b. As the iteration progresses, the simulations explored a broader conformational space and approached closer to the active state. We compared the performance of DASH to CMD (500 ns) and other non-CV-based enhanced sampling methods, including GaMD^29^ (500 ns) and replica-exchange with solute tempering^58^ (REST2, 200 ns × 40 replicas, 300-500K) (Fig.3c, Data S2, Section 2 of Supplementary Information). DASH outperformed all other methods in approaching the active state across all cases. To elucidate the superior performance of DASH, dimension reductions were performed to the trajectories using different CVs (CTMAE-generated CVs, RMSD-Rg and physical CVs by knowledge, see Supplementary Information S2). No matter in which CV space, DASH consistently covered a wider range of conformations than CMD (Fig.3d). In contrast, GaMD and REST2 achieved such expanded exploration in only 2-3 cases (demonstrated by markers). DASH trajectories reduced by each CVs were reweighted by MSM and the transition path as well as barriers were calculated (Supplementary Information S2). We found that barriers calculated from physical CVs are relatively low, with cases below 5k_B_T, probably because only parts of the protein motions are considered so that the difficulty of transition is underestimated. More importantly, cases that GaMD or REST2 performed relatively better (orange boxes in Fig.3e) have lower energy barriers, no matter calculated by which CVs. Meanwhile, in simulations of ADP-ribosylation factor 6^59^ (ARF6), although GaMD and REST2 sampled a larger conformational area, they failed to drive the system towards the active state (Fig. 3f), possibly due to their lack of direction predictions in their methodologies. The above results suggest that DASH’s enhanced performance stems from its combined ability of overcoming high energy barriers and orientation of FMS. Ablation studies confirmed the contributions of key DASH components (Fig. 3g), that seed hopping promoted broader conformational exploration, both CTMAE and seed selection help controlling the sampling direction. Additionally, removal of fragmentation hinders the sampling of the less flexible domain when multiple domains change during the state transition. The loss in performance would not be observed if only one domain is moved (Extended Data Fig. 2a).

**Fig. 3.**
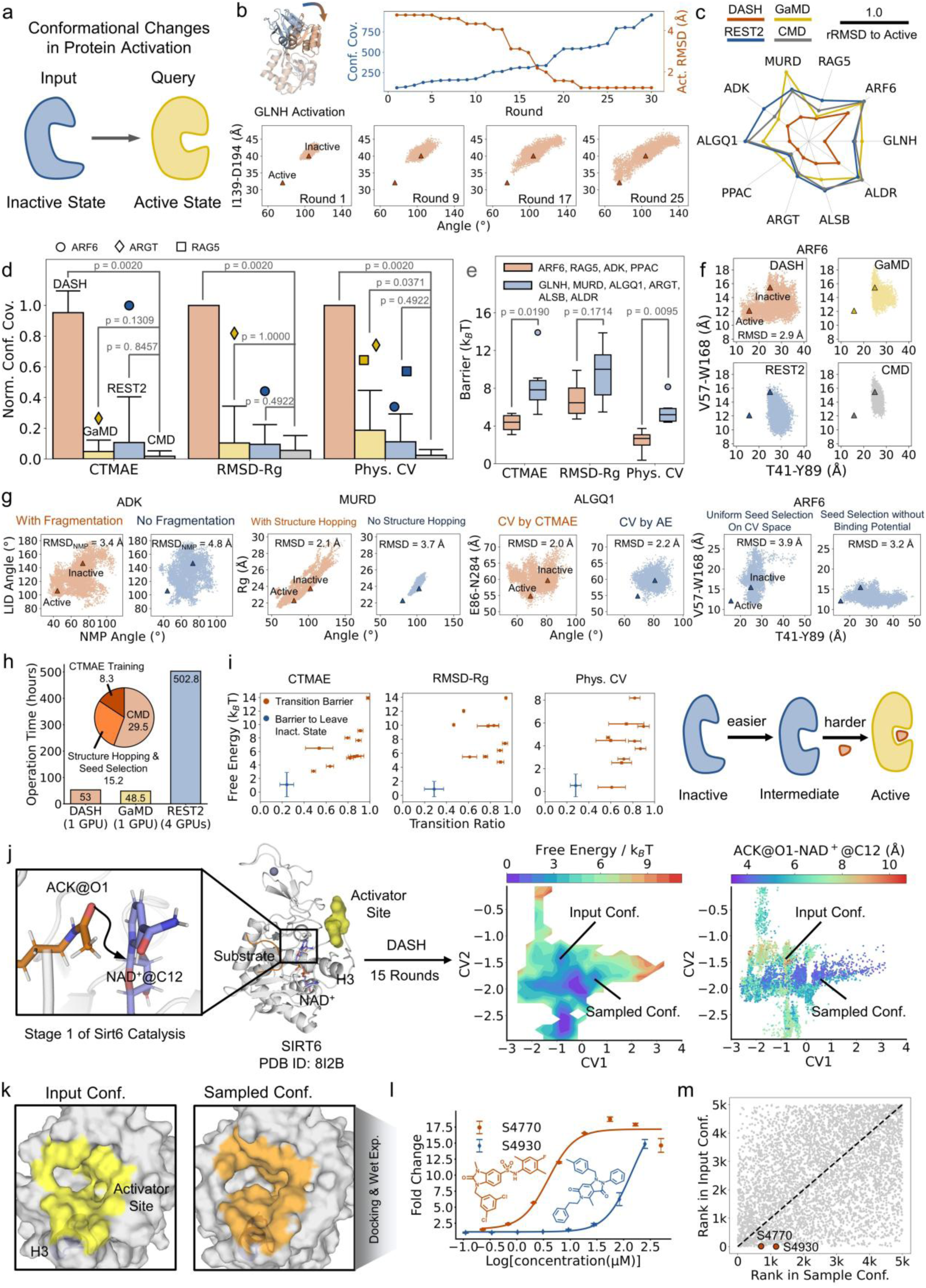
Sampling protein activation with DASH. **a.** Problem description and physical function for seed selection. **b.** An example of sampling the active conformation (orange) from inactive conformation (blue). Upper: change of conformation coverage RMSD to active state along iterations. Lower: change of ensemble along iterations. **c.** Comparison of minimum relative RMSD (rRMSD) to active state by different sampling methods (DASH, GaMD, REST2 and CMD). **d.** Comparison of normalized conformational coverage among different sampling methods. In brief, conformations were reduced to 2-D using collective variables (CVs), and the area in 2-D space is defined as the conformational coverage. Three types of CVs were used, including CTMAE-generated CVs, RMSD-Rg and physical CVs (case by case). Outliers were shown in markers, suggesting that GaMD or REST2 sample broader conformations in these cases. Circle: ARF6. Dimond: ARGT. Square: RAG5. Colors of the markers is the same to the corresponding sampling method. **e.** Comparison of transition barrier between cases that GaMD or REST performs better (orange) and worse (blue). Transition barriers were estimated on three types of CV-space. **f.** An example of larger conformational coverage but not boosted toward the active site in GaMD and REST2. **g.** Ablation studies. ADK: Removal of fragmentation. MURD: Removal of marginal structure hopping. ALGQ1: Use CTMAE or AutoEncoder for CV calculations. ARF6: Removal of seed selection. **h.** Bar plot: Operation time of sampling MURD by DASH (30 round × 10 ns, 1 RTX-3080 Ti), GaMD (500 ns, 1 RTX-3080 Ti) and REST2 (40 replicas × 200 ns, 4 RTX-3080 Ti). Pie plot: Time cost of each part in DASH. **i.** Orange: Average transition ratios and energy of conformations in transition state (TS) bin (see transition paths in Supplementary information and Methods section). The marker is square if the TS is at the end of the transition path and circle otherwise. Blue: Average transition ratio and energy for the bin whose transition ratio is closest to 0.25 on 10 cases. Right: A proposed mechanism of protein activation. Transition barriers were estimated on three types of CV-space. **j.** Sampling the active state conformation of SIRT6 from inactive state. Conformations are projected using CVs generated by CTMAE and reweighted by MSM. The contour plot (left) is colored by estimated free energy. The scatter plot is colored by the attack distance (see cartoon plot in the left) related to catalysis. **k.** comparison of the activator site in input conformation and the sampled conformation. **l.** Dose-dependent response of activators screened from the sampled conformation. **m.** Comparison of rankings in virtual screening using different conformations. Error bars represent standard deviations. Statistics analysis are analyzed using Student’s t-test (normal and homo-variance) or Mann-Whitney test (otherwise). N=10.

We compared the operation time among DASH, GaMD and REST2 on MurD with RTX-3090 Ti GPU. The operation time of DASH (53 hours) is comparable to that of GaMD (48.5 hours) and is only one-tenth of REST2 (502.8 hours) (Fig.3h). Among the 53 hours, 55% of the time was taken by CMD simulations, while NN training accounted for 16% of the time and hopping & selection accounted for the rest 29%. This reflects that structure hopping, namely SMD on NN-generated CVs is rather slow, which is only 21.6 ns per day. In comparison, the speed of CMD reaches 240 ns per day. The low speed might be explained by the low GPU utility (~20%) during SMD on NN-generated CVs. Therefore, in practice, seed hopping of multiple structures is performed in parallel on one GPU.

We analyzed the conformations at transition states (TS), which are calculated by different CV projections (CTMAE-generated CVs, RMSD-Rg and physical CVs). The conformations are measured by transition ratio, which is defined as the ratio of RMSDs towards inactive/active endpoint conformations. As shown in Fig. 3i, transition ratio of TS (orange) in most cases is above 0.5, suggesting that most TS resembled the active conformation. In some cases, the TS is virtually identical to the active conformation, indicating that the active conformation might be unstable without a ligand. We further calculated energy barrier to exit the inactive state, which is defined as conformations within a bin whose transition ratios are closest to 0.25. As a result, such barrier is relatively low (1-2 k_B_T). Collectively, these results suggest that escaping the inactive state during the inactive → active transition is facile. However, subsequent formation of active state is rate-limiting and usually necessitates ligand participation. This observation supports the classical induced-fit^60^ mechanism (Fig.3i).

Sirtuin 6 (SIRT6) is a NAD^+^-dependent lysine deacetylases that suppresses tumor progression^61^. Previous studies have shown that activator binding at an allosteric site near C-terminal^62–65^ would shift the conformation of H3 (60-66)^66, 67^, but whether such conformational changes related to activation of SIRT6 remains unknown. To answer this question, we performed a 15-round DASH on the inactive SIRT6 binding NAD^+^ and substrate. The resulting conformations were analyzed using CTMAE dimensionality reduction and MSM (Fig. 3j). SIRT6 catalysis initiates with a nucleophilic attack by the amide oxygen on the glucose carbon^66^. We found the attack distance correlates well with conformational changes of SIRT6, suggesting that DASH might have driven SIRT6 into a catalysis-favored conformation. Therefore, we predict the conformation near (0.5, −1.75) on the CV-space to be the potential active state, where most the attack distance is below 4.0 Å. The sampled conformation is formed by the inward-folding of H3 reshaping the activator binding site into a more concave pocket (Fig. 3k), according with previous MD analysis on activator-bound conformations^67^. We further screened molecules able to bind to activator site within the sampled conformation (Data S3). We purchased 10 compounds based on screening results (Extended Data Fig. 2b) for wet-lab validations, and in consequence we found two novel SIRT6 activator with EC_50_ of 3.9 μM and 114.1 μM and a maximum activation of 17.1 folds and roughly 20 folds, respectively (Fig. 3l, Table S2). The novel activators bind in a more concave position compared to previous activators (Extended Data Fig. 2b). Notably, we found the rankings of virtual screening is completely different when docking to the activator site in inactive conformation, with the discovered hits were ranked only at 708 and 1163. (Fig. 3m). In all, although many previous SIRT6 activators were discovered by its inactive conformation, our results demonstrate that identifying more functional conformations with DASH could help broaden the space of activator discovery.

### Sampling the opening of cryptic allosteric sites

Allosteric sites are druggable pockets on protein located differently from the orthosteric site^68, 69^. Designing modulators targeting allosteric sites are praised for better selectivity^70,71^. Many allosteric sites are open only in specific MSs and its stability requires binding of ligands, which are known as “cryptic allosteric sites”^72–74^. We tested whether DASH could sample FMS with cryptic allosteric site opening by testing on eight such cases (Fig. 4a, Section 3 of Supplementary Information). The DASH protocol was similar to that used for inactive → active state sampling, except only 25 rounds were used and smaller fragments were adopted in CTMAE, because pocket-opening usually change fewer residues. Two examples of DASH simulations, RORγt^5^ and HRAS^54^, are shown in Fig. 4b. As the iteration progresses, the cryptic allosteric site gradually opens. The pocket-open conformations formed a MS within the collective variable (CV) space generated by CTMAE (Fig.4c, Extended Data Fig. 3). We compared the performance of DASH with GaMD on the 8 cases. Unlike the task of sampling active state from the inactive state, GaMD can sample the open conformation in 7 of 8 cases. However, the degree of pocket exposure achieved by GaMD was generally less pronounced than that observed with DASH. These results validate the potential of DASH for discovering cryptic allosteric sites.

**Fig. 4.**
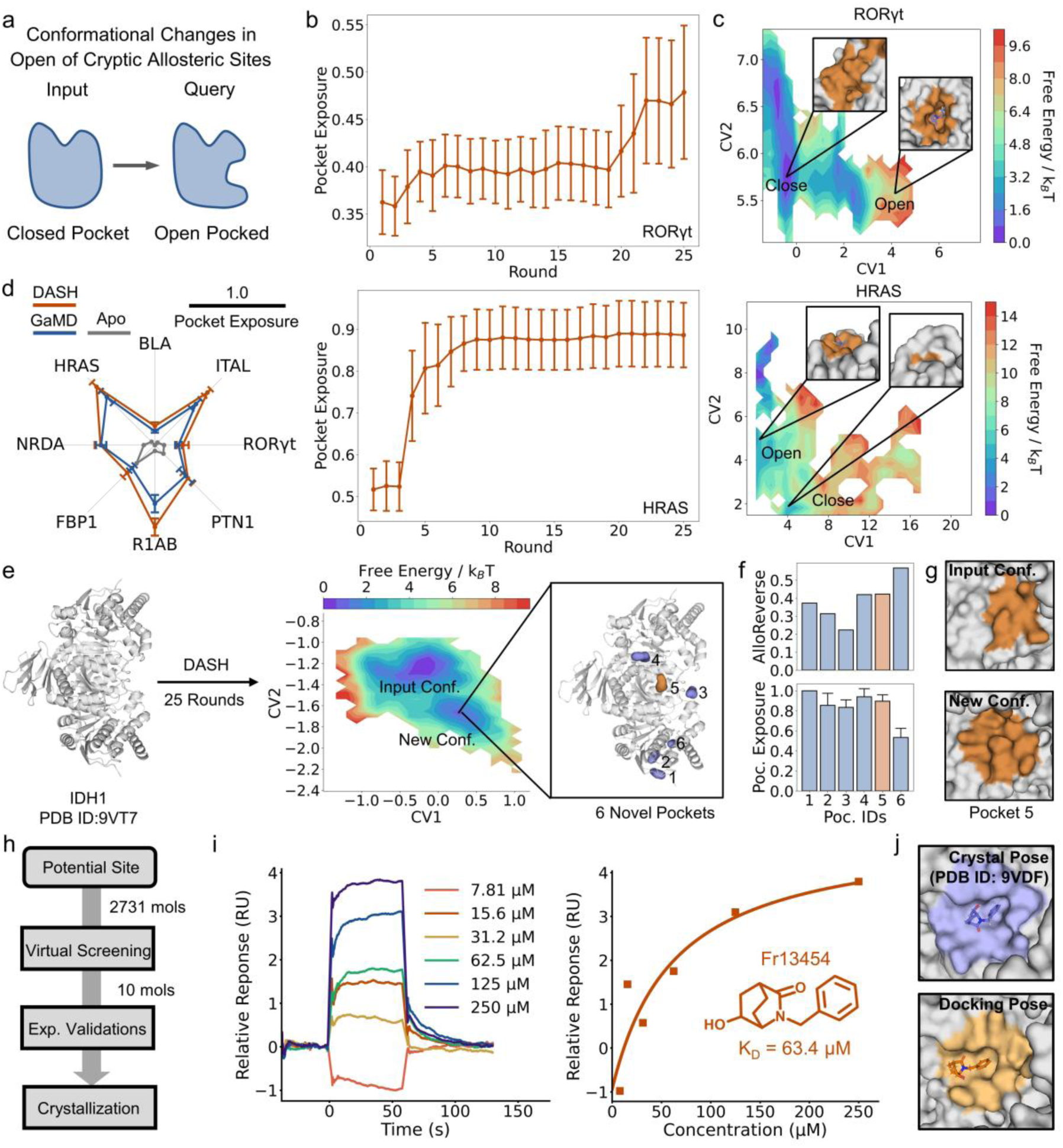
Sampling opening of cryptic allosteric site with DASH. **a.** Problem description and physical function for seed selection. **b-c.** Two examples of sampling the opening of cryptic allosteric site. **b.** Change of pocket exposure along iterations. Error bars represent standard deviations. **c.** Dimension reduction by CTMAE and metastable states for open/closed conformations. **d.** Comparison of pocket exposure by different MD methods. DASH is calculated using all rounds of trajectories. **e.** DASH simulation of *apo* IDH1, free energy surface (CTMAE CV reweighted by MSM), and novel pockets detected. **f.** Calculation of AlloReverse score and pocket exposure of the 6 pockets. **g.** Pocket 5 in different conformations. **h.** Validation workflow of pocket 5. **i.** SPR experiment of fragment Fr13454. **j.** Comparison of crystal and docking pose of Fr13454. Error bars represent standard deviations. N=400 for **b**, N=8 for **d**, N=10000 for **f** and N=3 for **i**.

IDH1 is a NADP^+^-dependent isocitrate dehydrogenase and is a famous drug target for glioblastoma multiforme and acute myeloid leukemia^75, 76^. To explore potential cryptic allosteric sites on IDH1, we performed a 25-round DASH on the *apo* IDH1. (Fig. 4e). A new conformation was observed, with 6 novel pockets detected by FPocket^77^. We employed AlloReverse^78^ score to evaluate their potential as allosteric sites and calculated their exposure along simulation. Collectively, Pocket 5 excel both at stability and AlloReverse score (Fig. 4f-g). To verify this predicted site, we conducted virtual screening and selected 10 molecules for wet-lab experiments (Fig. 4h, Extended Data Fig.4a, Fig. S3). Fr13454, a molecule with KD of 63.4 μM (Table S3), was further selected for co-crystallization with IDH1 (Fig. 4i, Table S1, Extended Data Fig.4b). The crystal structure confirmed that Fr13454 binds precisely within the predicted Pocket 5 (Fig. 4j), demonstrating the validity of the cryptic allosteric site identified by DASH. We noticed that the crystal pose of Fr13454 is more buried than the docking pose, suggesting that conformational deviations could exist between sampled pose and ligand-binding pose of cryptic allosteric sites. Together, these results show a successful example predicting novel cryptic allosteric site using DASH.

### Sampling helical folding of disordered regions

Although global or local movements of domains, namely cases in previous sections, account for most of conformational changes during protein functionalization, some proteins also undergo fold switching, like formation of secondary structures in intrinsic disordered regions (IDRs) (Fig. 5a). ^79^ However, DASH is not designed for sampling IDR folding. Fortunately, folding simulations has been well-studied using secondary-structure CVs (ssCVs)^80^. To address this gap, we integrated DASH with ssCVs. Briefly, during structure hopping, an additional biasing force is applied along relevant ssCVs to promote secondary structure formation (Fig. 5b). This hybrid approach, DASH+ssCV, might enable semi-directional sampling of fold-switching during NS→FMS transitions.

**Fig. 5.**
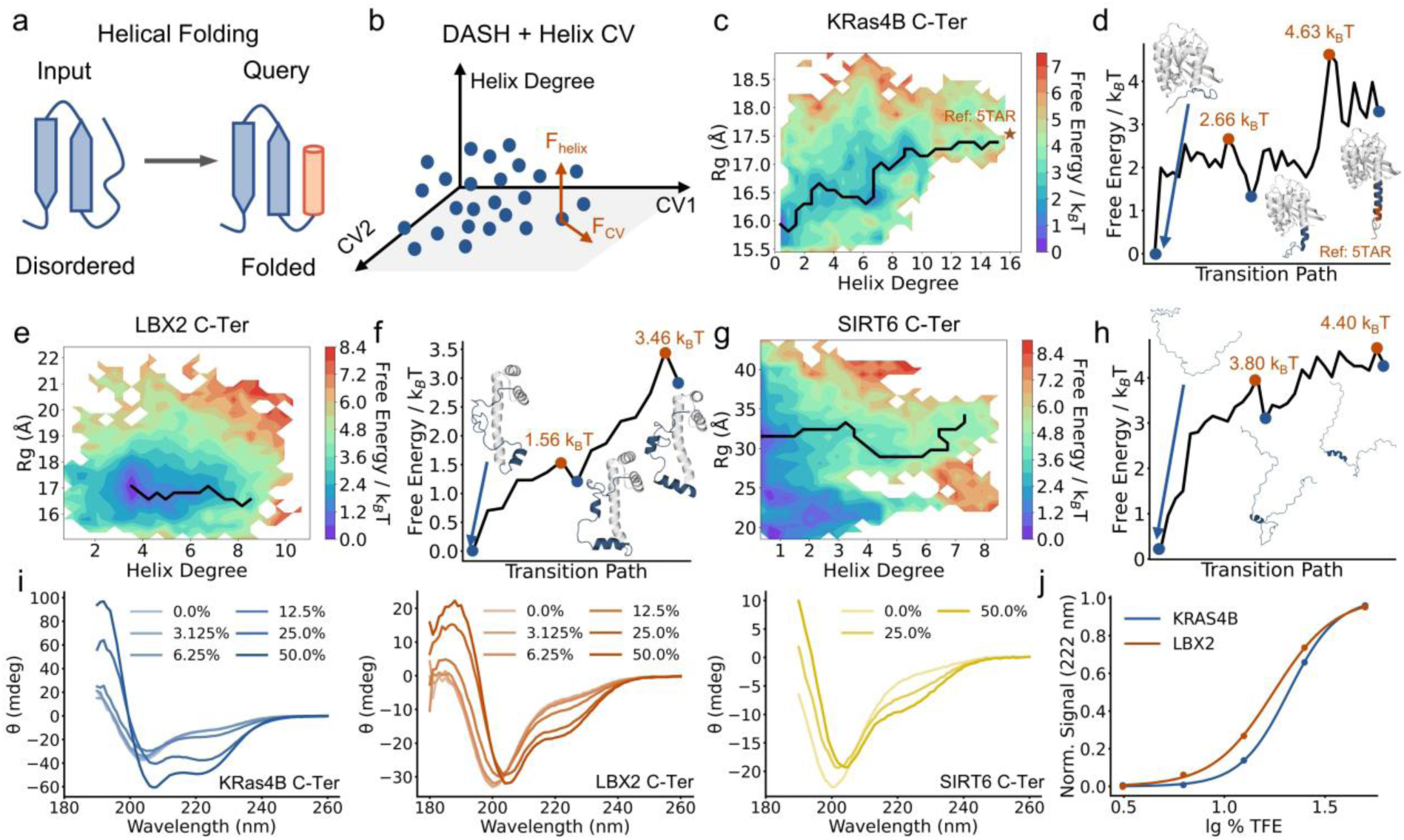
Sampling folding of disordered region by combining DASH and helical collective variables. **a.** Problem description. b. Illustration of method combination. A further force is applied on helix CV during seed hopping. **c,e,g.** Free energy surface of KRas4B C-terminal (**c**), LBX2 C-terminal (**e**) and SIRT6 C-terminal (**g**) sampled, all reweighted by MSM. **d,f,h.** Transition path of helix formation of KRas4B C-terminal (**d**), LBX2 C-terminal (**f**) and SIRT6 C-terminal (**h**). Local minima are labeled by blue dots and the representative structures are shown in cartoon. Transition states are labeled by orange dots and energies are shown. **i.** Circular dichroism spectrum of KRAS4B C-terminal, LBX2 C-terminal and SIRT6 C-Terminal with different ratio of trifluoroethanol (TFE). **j.** Relationship between normalized signal (helix degree) and ratio of TFE.

We tested this idea on KRas4B, an isoform of Ras protein, which plays critical roles in signal transductions related to cell growth and proliferation^81, 82^. It is also a star target in cancer treatment^83^. KRas4B has a disordered domain named “hypervariable region” (HVR, 165-185) in its C-terminal in its NS. HVR would favor a α-helical conformation when it interacts with PDE-δ during its transportation towards membrane^6^. We attempted to sample HVR folding using DASH+HelixCV^80^ from its NS^84, 85^. The 25-rounds simulation data (a total 250 ns) were projected using HelixCV (also helix degree) and radius of gyration (Rg). As shown in Fig. 5c, a folded HVR is found in a MS, with a small RMSD (2.2 Å) compared to reported folded state (PDB: 5TAR). The transition path from the free energy surface (FES) (Fig. 5d) suggested a step-wise folding mechanism, with two energy barriers of 2.66 k_B_T and 4.63 k_B_T. We noticed that either using HelixCV alone in well-tempered metadynamics (WT-MetaD)^86^ or DASH failed to sample the folded state (Extended Data Fig. 5a). Since folding of HVR contains both folding and domain swing, we speculate that WT-MetaD fail to overcome swing barrier while DASH fail to catch folding transitions. A combination of both DASH and HelixCV could tackle both obstacles.

We moved on to apply DASH+HelixCV on other IDRs. From the AlphaFold Database^87^, we found that the C-terminals of LBX2^88^ and SIRT6 are also disordered. IDPSAM^89^ were used to predict the C-terminals conformations of KRas4B, LBX2^88^ and SIRT6^90^, and it was found that the C-terminal of KRas4B and LBX2 could form α-helix while SIRT6 could not (Extended Data Fig. 5b-d). Motivated by the predictions, we performed 25-round DASH+HelixCV on LBX2 and SIRT6, and the trajectories were projected on helix degree and Rg (Fig. 5e-h). The transition paths indicate that the C-terminal folding of LBX2 and SIRT6 also adapt a step-wise mechanism. The FES reveals that C-terminal of SIRT6 populates disordered states more extensively than that of KRas4B and LBX2 and its folding involves a higher initial energy barrier (3.80 k_B_T) compared to KRas4B (2.66 k_B_T) and LBX2 (1.56 k_B_T). The computational findings correlate well with IDPSAM predictions, suggesting that helical formation is less favorable in SIRT6. Furthermore, LBX2 exhibits a lower folding barrier than KRas4B, suggesting its folding is easier than C-terminal of KRas4B, although the latter achieves a more fully folded conformation.

To validate these computational insights, we expressed the C-terminals of KRas4B, LBX2 and SIRT6 and induce formation of α-helix using trifluoroethanol^91^ (TFE). Helical formation was monitored by circular dichroism (CD) spectroscopy. Although in principle any IDR could be folded into α-helix with the help of TFE, their differentiated intrinsic power of helix folding make helix formations accessible under different concentrations of TFE^92–94^. In our study, KRas4B and LBX2 show a dose-dependent formation of α-helix (Fig. 5i, Data S4) with low concentration of TFE, whereas SIRT6 showed only a weak helical signal even at high TFE concentration, consistent with its predicted folding difficulty. The stronger CD signal for KRas4B compared to LBX2 suggests a greater proportion of folded residues, corroborating our folding states sampled (Fig. 5c, e). The half-formation-concentration of LBX2 (17.6% TFE) is smaller than that of KRas4B (20.8% TFE) (Fig. 5j), in agreement with its lower predicted transition barrier (Fig. 5d, f). Collectively, the above results demonstrate a successful example to predict potent helix-formation in disordered regions with DASH+HelixCV.

## Discussion

Identifying function-related metastable states (FMS) and its transition pathway from only a given native state (NS) is an attractive topic in biology and pharmacy^7–12^. However, molecular dynamics (MD) sampling FMS from NS is underexplored, possible because this is an energy-ascending process^42^. CMD simulations of such processes, though unbiased and more reliable, usually require careful selection of several starting conformations^95, 96^ or introducing propriate mutations^97^, which is case-by-case and does not have a general tactic. Collective-variable-based (CV-based)^23–25^ and transition-pathway-based^26–28^ methods are hard to be applied for the difficulty of assigning sampling directions when only the initial state (NS) is known, while methods require no direction assignments, as our results demonstrate, either fail to overcome the high energy barrier or inadvertently drive the system towards irrelevant states. To realize sampling of FMS with only NS known, we introduced an adaptive sampling method named DASH. DASH utilizes a combined contrastive learning and masked autoencoder (CTMAE) neural network to analyze accumulated trajectories and yield CVs presenting tendencies of conformations transitions. Subsequent structure hopping and seed selection are used to selectively amplify tendencies more likely towards FMS.

We first tested DASH on two kinds of NS → FMS tasks, including 10 cases of protein activation and 8 cases of open of cryptic allosteric sites. DASH succeeded in all cases, significantly outperforming other enhanced sampling methods without explicit direction assignment (GaMD and REST2) and require comparable time to GaMD. We further combined DASH with helical collective variables to reemerge helix-formation of KRas4B HVR. More importantly, we applied DASH to sample an active conformation of SIRT6, facilitating the screening and discovery of a novel activator, and we discovered a novel cryptic allosteric site on IDH1 with a hit binding to it. Since activators and cryptic allosteric sites are usually hard to discovered in NS, DASH has shown the potential to enlarge the druggable space. Besides, targeting IDR^6^ has drawn great attentions nowadays. DASH combined with directional methods (DASH+ssCV, and mainly DASH+HelixCV here) has also been applied to found a new foldable IDR in LBX2.

DASH, adaptive sampling^33, 34^ (AS) and adaptive CV-based methods^35–41^ (ACV) all operate in an iterative workflow, that analyze accumulated trajectories to guide new rounds of MD simulation. AS identifies lack-sampled regions from trajectories and starts CMD from them. Although it is unbiased, there is no guarantee that the new simulations would overcome transition barriers. Comparatively, ACV generates transition-related CVs from previous simulations, usually via neural networks (NNs), and starts new biased simulations along the CVs so that the conformations on the directions would be well-sampled and relative transition barriers would be overcome. However, as shown in Fig. 3h, calculating NN-based CVs is very time-consuming. Our results shows that the speed of MurD simulation is 10-times shrunk from 240 ns/day to 21.6 ns/day when using NN-CVs. DASH combined the advantages of both AS and ACV. It generates CVs like ACV, but only applies CVs to run short biased simulations to help marginal conformation to across transition barriers, namely the “structure hopping”. Selected hopped structures are used as seeds to start new CMD simulation, just like AS. Another important innovation of DASH is seed selection, which narrow the sampling directions into those more likely towards FMS via physical filters. Previous AS and AVC could drive the system into any conformations rather than targeting FMS.

In conclusion, we have developed DASH, a method capable of efficiently sampling FMS conformations starting from NS only. DASH have been extensively benchmarked and utilized for discovery of new conformations, new binding sites, and new activators. Since boosting directions in DASH are learned from previous trajectories, conformational sampling of rigid regions or complex movements like IDR folding is difficult with DASH alone. Future work could focus on making DASH a more general sampling method.

## Methods

### Simulation setup and conventional molecular dynamics (CMD)

Most the simulation topology and parameter files were prepared using *tleap* in AMBER20^98^ suite. Missing loops were amended using PDBFixer^99^. Missing atoms are added, and solvation box were built with TIP3P water (12 Å shell thickness). Systems are neutralized by Na^+^ and Cl^-^. The protein is described with ff14SB forcefield^100^. Ligand parameters are either obtained from AMBER parameter database (http://amber.manchester.ac.uk/) or GAFF2 forcefield with AM1-BCC^101, 102^ derived charges. Zinc ions, coordinated by neighboring serine residues, were described using the ZAFF forcefield^103^. C-terminal of SIRT6 was prepared in the same simulation condition but using CHARMM36m^104^ forcefield using CHARMMGUI^105^.

CMD in DASH, including initial simulation and seeded simulations, are all performed by OpenMM^106^, while the rest CMD simulations were performed by GPU-accelerated pmemd in AMBER20^98^. In general, the systems are minimized, followed a gradual heating and equilibration. The productions runs are conducted in NPT ensemble at 300K, employing a Langevin thermostat with a collision frequency of 1 ps^-1^. The integration step is 2 FMS and the pressure is maintained at 1 bar.

### Contrastive learning and masked autoencoder (CTMAE)

As shown in Fig. 1c, we defined ***c***_***i***_ to be the i^th^ frame of the input Cα trajectory and ***x***_***i***_ to be it flattened version, namely a *N*_*atom*_ × 3 dimention vector. To define CTMAE neural networks (NNs), fragmentation is first performed. Root mean square fluctuation (RMSF) is calculated on {***c***_***i***_}. Residues within continuous regions exhibiting RMSF values above a cutoff are selected and clustered into two fragments using KMeans^107^ based on their coordinates in the initial structure. The minimum size for a continuous region is 5 residues, reduced to 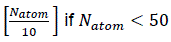. The cutoff is typically the mean RMSF, but set to 0 if *N*_*atom*_ < 20. For sampling cryptic allosteric site opening, clustering is replaced by selection using a scoring function that evaluates the linear correlation between trajectories of continuous regions and the full trajectory {***c***_***i***_}. This process yields two fragment trajectories with *N*_1_ and *N*_2_ atoms, and their flattened vectors of the i^th^ frame are denoted as 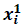 and 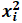.

Based on the defined fragments, we define two multi-layer perceptron (MLP) encoders with LeakyReLU active functions (negative slope set at 0.01) with layer dimensions of [*N*_1_ × 3] × [*N*_1_ × 3] × 8 × 1 and [*N*_2_ × 3] × [*N*_2_ × 3] × 8 × 1, to map {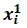} and {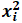} to two-dimensional collective variables (CVs) {***z***_***i***_}. Graph encoders were also tested (see Supplementary Information). To train the encoder, we further define a scorer (2 × 8 × 1) and a decoder (2× 8 × [*N*_*atom*_ × 3] × [*N*_*atom*_ × 3]). The scorer takes pairwise differences (***z***_***i***_ − ***z***_***j***_)^2^ as input and regress the root mean square distance (RMSD) between ***c***_***i***_ and ***c***_***j***_, while the decoder reconstructs ***x***_***i***_ from ***z***_***i***_. The total loss is the weighted sum of regression loss and reconstruction loss. Model construction and training were implemented in PyTorch^108^.

### Structure hopping

CV space is discretized into grids. Marginal grids are identified and grouped into 20 clusters using KMeans. The cluster centers are denoted as {***m***_***i***_}. Marginal conformations are selected as the closest frames of each ***m***_***i***_. For the i^th^ marginal conformational, a step-wise steered MD simulation^55^ is performed. The simulation setup is generally the same as CMD, except an additional harmonic force is applied. For the j^th^ round,

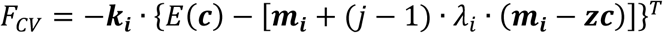

where ***k***_***i***_ is a vector of harmonic force constants. *E* is the trained encoder converting coordinates aligned to input structure into CVs. ***zc*** is the center of CV space. The term ***m***_***i***_ + (*j* − 1) · *λ*_*i*_ · (***m***_***i***_ − ***zc***) is a stepwise-updated harmonic center to steer the system away from ***zc***. *λ*_*i*_ is a scaling factor controlling a balanced hopping among different marginal conformations. Each round of seed hopping contains a 10 ps simulation, realized by OpenMM^106^. CVs are calculated along simulation with openmm-torch package^106^. Seed hopping is stopped when the maximum RMSD between hopped structures and previous trajectories is above a cutoff value, which is 20% percentile of pairwise RMSD in trajectory and capped at 2 Å.

### Seed selection

Potential energy of each successfully hopped seeds are calculated as the average potential energy in the last 2 ps hopping simulation (recorded every 40 fs). To calculate the binding potential, solvents and hydrogens are removed from the final hopped structure, followed by pocket detection with AutoSite^109^. The binding potential is defined as the average pocket energy of top 50% identified pockets. Hopped seeds are first sorted by the ascending of potential energy. For folding disordered regions, the top 4 seeds are selected. For other applications, the top 8 seeds are re-ranked by binding potential to select the final 4 seeds.

Seed selection by SASA and burriedness are done by first detecting pockets on protein surface with FPocket4, which better finding cavities because it is a geometric method. Pocket SASA is summed to be SASA and pocket volume is summed to be burriedness.

### Markov state modeling (MSM), free energy surface, and transition path

MSMs are constructed using the pyEMMA^44^ package. CV space data are first clustered into 50 microstates, and MSM is built on these microstates with a lag time of 0.5 ns. Estimation of implied timescales suggests that a lag time of 0.5 ns has reached a convergence in most cases (see Supplementary Information section 2-4). MSM were further validated using Chapman-Kolmogorov tests (see Supplementary Information section 2-4), bootstrapping analysis and constructed with longer lag time (Supplementary Information section 5, Figure S1-S2), and the results of energy barriers were not changed by adjusting these parameters. Based on the constructed MSM, the CV space is discretized into grids and reweighted. Free energy is calculated on each grid using Boltzmann distribution. Native state (NS) and function-related metastable state (FMS) are assigned to grids, and the path connecting the two girds with minimal energy barrier is calculated with Dijkstra algorithm, yielding the transition barrier energy.

### Benchmarking dataset of CTMAE

We collected 15 trajectories from D.E.Shaw Research and our previous published work of MD. The dataset contains 8 fast-folding proteins^110^ (chignolin, Trp-cage, BBA, Villin, WW domain, NTL9, Protein B and α3D) and 7 globular proteins (AKR^111^, SIRT6^61^, NDR1 (not published), EGFR^112^, KRas4B^85^, ACE2^113^ and 2NMBR^114^).

### Dimension reduction methods for comparison

We compared CTMAE with principal component analysis^48^ (PCA), TICA^50^, IsoMap^49^, UMap^115^, Autoencoder^35^ and EncoderMap^116^. To align with CTMAE, the encoder structure is a NN [*N*_*atom*_ × 3] × [*N*_*atom*_ × 3] × 16 × 2, except for PCA, which use the matrix after decomposition as encoder. The differences mainly lie in loss functions, which compare, the deviations of pairwise distances between high-D and low-D space (IsoMap), consistency of neighboring relationship between high-D and low-D space^117^ (UMap), reconstruction loss (AutoEnoder) and reconstruction loss plus specific distance relationships between high-D and low-D space (EncoderMap).

### Micro-/Macro-state consistency

This metric evaluates whether conformations within small CV regions are structurally similar, indicating thermodynamic preservation. CV space is discretized into 40×40 (micro-state) and 10×10 (macro-state) grids. Maximum pairwise RMSD is calculated for each grid, if possible. Average RMSDs are taken as consistency. For cross-method/case comparison, values were normalized per case:

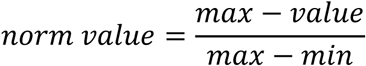

so that the method with the best consistency will be scored 1.

### Recovery of secondary structure

This metric also reflects thermodynamic preservation, calculated for chignolin, Trp-cage, Villin, WW domain, Protein B, and KRas4B. The secondary structure of a frame is defined as a vector, where each dimension represents helix/sheet degree of a folding area. The helix/sheet degree follows previous definition^80^:

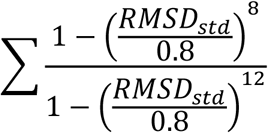

where *RMSD*_*std*_ is the RMSD between a six-residue fragment and a standard helix or sheet (parallel or antiparallel) template. We use normalized CV values as input to regress the secondary structure vector with a MLP NN (2×4×8×4×1). The recovery is defined as the coefficient of determination of the MLP regression. The training and scoring are realized using Sklearn^118^ toolkit. Values are normalized per case:

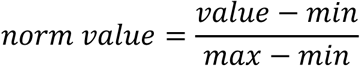

so that the method with the best recovery of secondary structure will be scored 1.

### Margin to all / intra margin RMSD

This metric reflects the diversity of marginal structures, which ensures a more diverse sampling in following iterations. Marginal structures are generated following “**Seed hopping**” section. RMSDs are calculated between the marginal conformation to the whole trajectory (Margin to All RMSD) or other marginal conformations (Intra Margin RMSD). RMSDs are normalized as previous section, so that method with the highest diversity (highest RMSD) scores 1.

### Dimension reduction algorithms for DASH generated data

To compare DASH with other methods in sampling of protein activation, we performed dimension reduction with different CVs, including CTMAE-generated CVs from DASH trajectories, RMSD combined with radius of gyration (Rg), and intuitive physical CVs. based on inactive-to-active transitions (listed in Supplementary Information).

### Relative RMSD (rRMSD)

To compare the approaching of active state from inactive state in different cases, we calculated rRMSD to be the ratio of minimum RMSD to active state and the RMSD between inactive/active state.

### Conformational coverage

This score indicates the explored conformational range. Normalized CV space dots were used to calculate the covered area via the convex hull method implemented in SciPy. To compare conformational coverage in different cases, the value is normalized per case using the formula in “**Recovery of secondary structure**”, so that method exploring the largest range of conformation will be scored 1.

### Gaussian-accelerated molecular dynamics (GaMD)

GaMD^29^ is performed using GPU-accelerated pmemd in AMBER20 package. The preparation step is the same as CMD. After minimization, a 10 ns CMD is carried to collect the initial boost potential and this potential is continuously updated in a following 50 ns biased simulation. The boost potential is added on both potential energy and dihedrals. Biased simulation on fixed boost potential is used for 500 ns GaMD production.

### Replica exchange with solute tempering (REST2)

REST2^58^ is performed using OpenMM. 40 replicas with temperature from 300 K to 500 K are run in parallel. All replicas are first equilibrated for 50 ps. Exchange attempts occurred every 1 ps. Simulations were limited to 200 ns per case due to extensive computational cost.

### Transition ratio

Transition ratio is a metric to evaluation the position of a conformation on inactive → active transition, based on the similarity compared to the two states. The transition ratio of the i^th^ frame of the trajectory is:

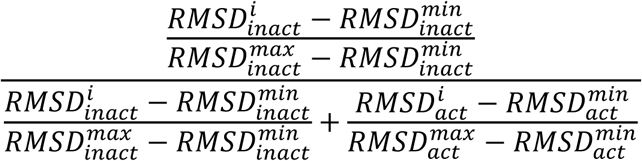

where 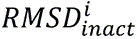 and 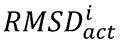 is the RMSD towards inactive and active state, respectively. 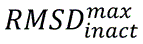, 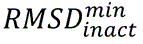 are the minimum and maximum RMSD value in the trajectory towards inactive state whereas 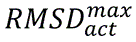, 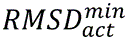 are the minimum and maximum RMSD value in the trajectory towards active state.

### Virtual screening

Molecular dockings were all performed with Schrödinger suite. Proteins are prepared and minimized using Protein Preparation Wizard. The docking box is defined from a given center, with an inner cubic box of 6 Å and an outer cubic box of 26 Å. Dockings were performed with Glide^119, 120^. The library for virtual screening is L7800 (https://www.targetmol.com/compound-library/High_Solubility_Fragment_Library) for SIRT6 and a manual library (Data S3) for IDH1.

### Protein expression and purification of SIRT6

The wild-type SIRT6 protein was constructed by direct insertion into the pET28a-His vector, which was transformed into commercial receptive Escherichia coli Rosetta (DE3) and cultured overnight on LB plates resistant to kanamycin. Single colonies were selected and cultured in 25 mL LB liquid medium with kanamycin resistance. After overnight culture, the cells were transferred to 1 L medium and cultured at 37℃, until the OD600 value was about 0.6-0.8. After 0.5 mmol/L IPTG was added, the cells were induced at 16℃ for 16-18h, collected by centrifugation at 4℃, resuspended with lysis buffer (300 mmol/L NaCl; 5% glycerol; 1×PBS; adjust the pH to 7.5), 1 mmol/L DTT and Protease Inhibitor Cocktail (K1012, APExBIO) in proportion. The wild-type SIRT6 protein was purified by Ni^2+^ column affinity chromatography and then centrifuged with SIRT6 assay buffer (50 mmol/L Tris-HCl; 137 mmol/L NaCl; 2.7 mmol/L KCl; 1 mmol/L MgCl_2_, pH adjusted to 8.0) several times to replace the elution buffer.

### FDL assays of SIRT6

The FDL assay was performed as described previously^63^. All compounds were dissolved with DMSO and diluted to the concentrations of 1 mmol/L. The 50 μL reaction system contained 5 mmol/L SIRT6, 2.5 mmol/L NAD^+^, 75 mmol/L RHKK-ac-AMC, 5 μL compounds/DMSO, and assay buffer. The reactions were conducted at 37 ℃ for 2.5 h, terminated with 40 mmol/L nicotinamide, and developed with 6 mg/mL trypsin for 30 min at 25℃. FDL assays for each compound were independently repeated at least three times.

### Pocket exposure

This metric evaluates cryptic site opening, defined as the recall of pocket residues present in the *holo* conformation. First, FPocket 4.0^77^ detected pockets in *holo* structures. The pocket best aligning with the allosteric ligand was selected and its residues are recorded as pocket residues. For BLA (PDB ID: 1PZP), since a too large pocket is predicted by FPocket, pocket residues are manually defined as those within 6 Å of the terminal phenylamino group. Then, we perform FPocket detection on each frame of trajectory. FPocket then analyzed each trajectory frame. For each frame, the highest residue recall among detected pockets defined its exposure. We then define the top 10% exposure values of accumulated frames to be the exposure of that round.

### AlloReverse

AlloReverse^78^ evaluates the potential of a Fpocket-detected pocket to be an allosteric site. It utilizes AdaBoost with input including mean local hydrophobic density, flexibility and reversed allosteric effect (RAE) as input. The first two descriptors are fetched from FPocket while RAE is the change of pairwise energy in pocket calculated via normal mode analysis. Inputs were z-score normalized. Higher AlloReverse scores indicate greater allosteric potential.

### Cloning, overexpression, and purification of IDH1

The IDH1 gene was cloned into the pET21b vector and transformed into Escherichia coli BL21 (DE3) competent cells. Positive colonies were selected on LB agar plates containing ampicillin and cultured in liquid LB medium. Protein overexpression was induced by adding 0.5 mM IPTG when the optical density at 600 nm (OD600) reached 0.6–0.8, followed by incubation overnight at 18°C. After incubation, bacterial cells were harvested by centrifugation at 4000 rpm for 10 min and resuspended in lysis buffer (150 mM NaCl, 25 mM Tris-HCl, pH 8.0). Cells were lysed using ultrasonication, and the lysate was centrifuged at 13,000 rpm for 20 min at 4°C to remove cell debris. The clarified supernatant was applied to Ni-NTA affinity chromatography resin, and IDH1 protein was eluted using lysis buffer containing 300 mM imidazole. Further purification was performed by size-exclusion chromatography (SEC) using a HiLoad 26/600 Superdex 200 pg column (GE Healthcare, UK), equilibrated with lysis buffer. Peak fractions containing purified IDH1 protein were collected, concentrated to approximately 10 mg/mL, aliquoted into 100 µL portions per tube, flash-frozen in liquid nitrogen, and stored at −80°C for subsequent crystallization experiments.

### Surface plasmon resonance (SPR) of IDH1

SPR experiments were performed on a Biacore 8K instrument (Cytiva). Full length IDH1, with a N-terminal 6×His tag, was immobilized on a CM5 sensor chip at a concentration of 50 μg/mL in 10 mM sodium acetate coupling buffer (pH 5.0) to a surface density of 11,000 Response Units (RU) by amine coupling chemistry. The surface was first activated with a 1:1 mixture of EDC/NHS (contact time: 420 s, flow rate: 10 μL/min), followed by the immobilization of IDH1, then deactivated with 1 M ethanolamine (pH 8.5). Binding experiments were performed at 25 °C using a running buffer of 10 mM Na2HPO4, 1.8 mM KH2PO4, 137 mM NaCl, 2.7 mM KCl, 0.05% Tween 20, 2% DMSO, pH 7.4. Compounds were 2-fold serially diluted using the running buffer and injected over the sensor surface at a flow rate of 30 μL/min at 25 °C with an association time of 60 s and a dissociation time of 60 s. Solvent corrections were performed using running buffers containing 1.6%–2.4% DMSO. Data were fitted with steady-state affinity analysis using Biacore 8K evaluation software.

### Crystallization and small molecule soaking

Crystallization of the IDH1 protein was carried out at 4°C using the hanging drop vapor diffusion method. The crystallization condition consisted of 0.2 M Li_2_SO_4_, 0.1 M HEPES (pH=7.4), and 20% PEG6000. Crystals appear in two days. For the small molecule soaking procedure, 0.5 µL of the fragment compound with 100 mM was mixed with 0.5 µL of reservoir solution, and the mixture was then added to the crystallization well containing the crystal. The final concentration of the small molecules was approximately 20 mM. The soaking duration was set to at least 24 hours. After soaking, all crystals were immediately flash-frozen in liquid nitrogen and stored after being treated with a cryoprotectant solution containing 25% glycerol and crystallization buffer.

### Structure determination

X-ray diffraction data were collected at beamline BL19U1 of the Shanghai Synchrotron Radiation Facility (SSRF) and processed using XDS^121, 122^. Data merging and scaling were performed using Aimless from the CCP4 suite^123^. The crystal structure was solved by molecular replacement using Phaser, employing the previously reported IDH1 structure (PDB ID: 5SVO) as the search model^124, 125^. The model was iteratively refined and built with Phenix Refinement and Coot, leading to the final structure^126^. The data collection and refinement statistics are summarized in Table S1.

### DASH combined with helix CV

Helix CV is exactly the helix degree defined in “Recovery of secondary structure”. The combined scheme generally follows DASH scheme. The only difference lies in structure hopping. An additional force is added during the SMD in structure hopping to encourage formation of α-helix. As shown below:

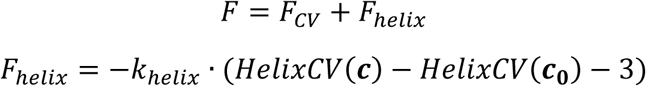

where *F*_*CV*_is the hopping force defined in DASH, *F*_*helix*_is the force to promote helix formation. *HelixCV* is the function to calculate helix degree. ***c*** is the coordinate of protein and ***c***_**0**_ is the initial coordinate of the marginal structure to be hopped. *k*_*helix*_ is the harmonic force constant.

### Well-tempered metadynamics

The well-tempered metadynamics (WT-MetaD)^86^ of KRas4B is performed using OpenMM^106^, using helix degree and radius of gyration (Rg) as CVs. The simulation setup is generally the same with CMD, except an accumulated biased potential is added to the system. Biased potential in WT-MetaD is calculated by grid method. The helix CV is calculated for residue from 160 to 180, so that it range is set from 0 to 16, with a bias width of 0.8. Since the Rg of IDR state and helix state KRas4B is 15.7 Å and 17.5 Å, we set the range of Rg CV from 15.0 Å to 20.0 Å, with a bias width of 0.5 Å. The bias factor is set at 4.0 while the initial height of Gaussian is set at 10 KJ/mol. The biased potential is updated every 2 ps.

### Protein expression of IDRs

Recombinant pET28a plasmids, bearing a N-terminal His×6 tag followed by a SUMO tag, were constructed to express the C-terminus of LBX2 (UniProt entry: Q6XYB7, residues 157–198, with a terminal tryptophan for UV monitoring), KRas4B (UniProt entry: P01116-2, residues 164–188), or SIRT6 (UniProt entry: Q8N6T7, residues 271 to 355). The proteins were overexpressed in E. coli Rosetta (DE3) grown in 2×YT medium induced by 0.3 mM IPTG in 16 hours at 289K. After cell disruption using high-pressure homogenization and centrifugation of the lysates, the proteins were initially purified using a nickel column and subsequently cleaved by ULP1 in 6 hours. After the second nickel column purification to remove the two tags, the proteins were further purified by size exclusion chromatography on a Superdex-30 column using an ÄKTA FPLC system (GE Healthcare). The purified proteins were dialyzed into phosphate-buffered saline (PBS, pH 7.4) and concentrated to approximately 10.0 mg/mL.

### Conformational exchange assay using circular dichroism spectroscopy

All circular dichroism (CD) spectra were acquired on a Chirascan VX spectropolarimeter (Applied Photophysics Ltd., UK) equipped with a Peltier-type temperature controller set at 298 K. Each spectrum was the average of three consecutive scans, with the buffer serving as the blank. Far-UV CD spectra were collected in the wavelength range of 190–260 nm using a 1-mm path length cuvette. The samples contained 0.1 mg/mL protein in PBS (pH 7.4) or varying concentrations of 2,2,2-trifluoroethanol (TFE). The spectra were recorded in continuous scanning mode at a rate of 50 nm/min with a bandwidth of 1 nm. Data analysis was performed using Pro-Data Viewer software integrated with the spectropolarimeter.

## Data Availability

All protein structures were downloaded from Protein Data Bank (https://www.rcsb.org/). Library for screening SIRT6 activators were downloaded from https://www.targetmol.com/compound-library/High_Solubility_Fragment_Library. Input files for DASH simulations in this article could be found in out GitHub page (see Code Availability). Solved crystal structures have been uploaded to Protein Data Bank, with accession codes of 9VT7 (*apo* IDH1) and 9VDF (IDH1-Fr13454 complex). MD trajectories for dimension reduction in this article are available at https://zenodo.org/records/16917262.

## Code Availability

Codes for DASH sampling and dimension reduction are available at https://github.com/JinyinZha/DASH. Tutorials of DASH could also be found in the website.

## Acknowledgements

This study was partly supported by grants from the National Key R&D program of China (2023YFF1205103 to J.Zhang), the National Natural Science Foundation of China (81925034, 82441035, 22237005 to J.Zhang, 32300531 to X.L.), Innovative research team of high-level local universities in Shanghai (SHSMU-ZDCX20212700 to J.Zhang), the Starry Night Science Fund of Zhejiang University Shanghai Institute for Advanced Study (SN-ZJU-SIAS-007 to J.Zhang.), the Key Research and Development Program of Ningxia Hui Autonomous Region (2022BEG01002 to J.Zhang), Shanghai Municipal Health Commission (2025ZHYL038 to J. Zhang), and Lingang Laboratory (LG8888 to J. Zhang). We thank D.E. Shaw Research for providing molecular dynamics trajectories.

## Author Contributions

J.Zhang conceptualized the study. J.Zha developed the DASH algorithms. Z.Z. performed experiments of disordered proteins. J.Zhong performed SPR experiments of IDH1. W.W., Q.X. and W.Q. solved the crystal structures of IDH1. Q.L., Q.S. and L.F. performed experiments of SIRT6. M.L., C.W., Q.R., N.L., H.Z. and X.L. generated the trajectories and data analyzed in the study. J.Zhang, L.F. and W.Q. supervised the study. J.Zha, Z.Z., J.Zhong, Q.Li and W.W. wrote the original draft. All the authors reviewed and edited the article.

## Competing Interests

The authors declare no competing interests.

---

**Extended Data Fig. 1.**
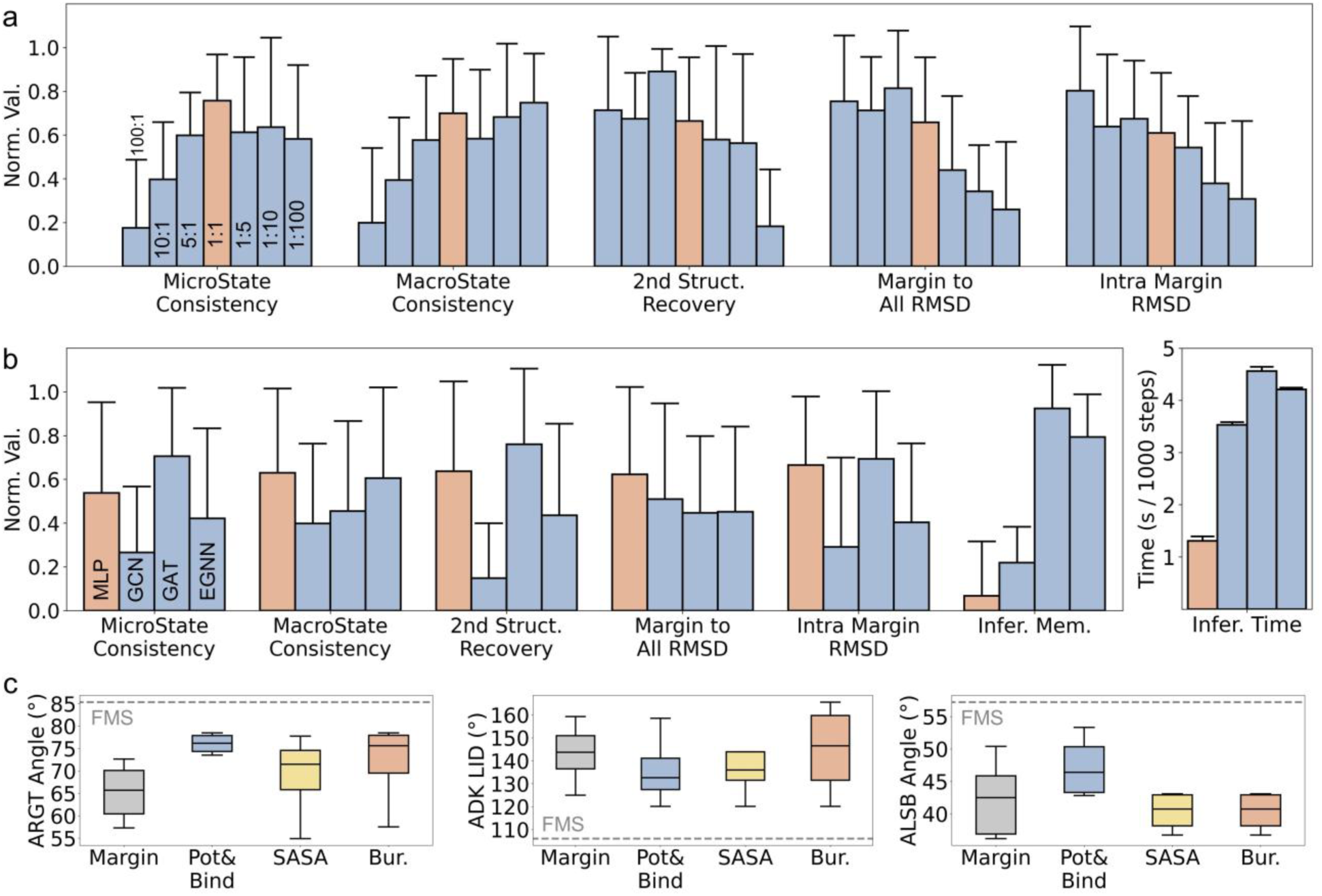
Performance of CTMAE under different circumstances. **a.** Ratio of coefficients of the contrastive (CT) loss and masked autoencoder (MAE) loss. **b.** Types of CTMAE encoders. N=15. Error bars represent standard deviations. **c.** Effect of different seed selection filters in sampling function-related metastable state (FMS) conformations. Margin: marginal conformations. Pot&Bind: potential energy and ligand-binding potential (our chosen methods); Bur: burriedness. N=4. Box plot represents the range between 25% and 75% quantile, with line on it represents the median value. Whiskers of box plot represents 1.5 times of inter-quantile range.

**Extended Data Fig. 2.**
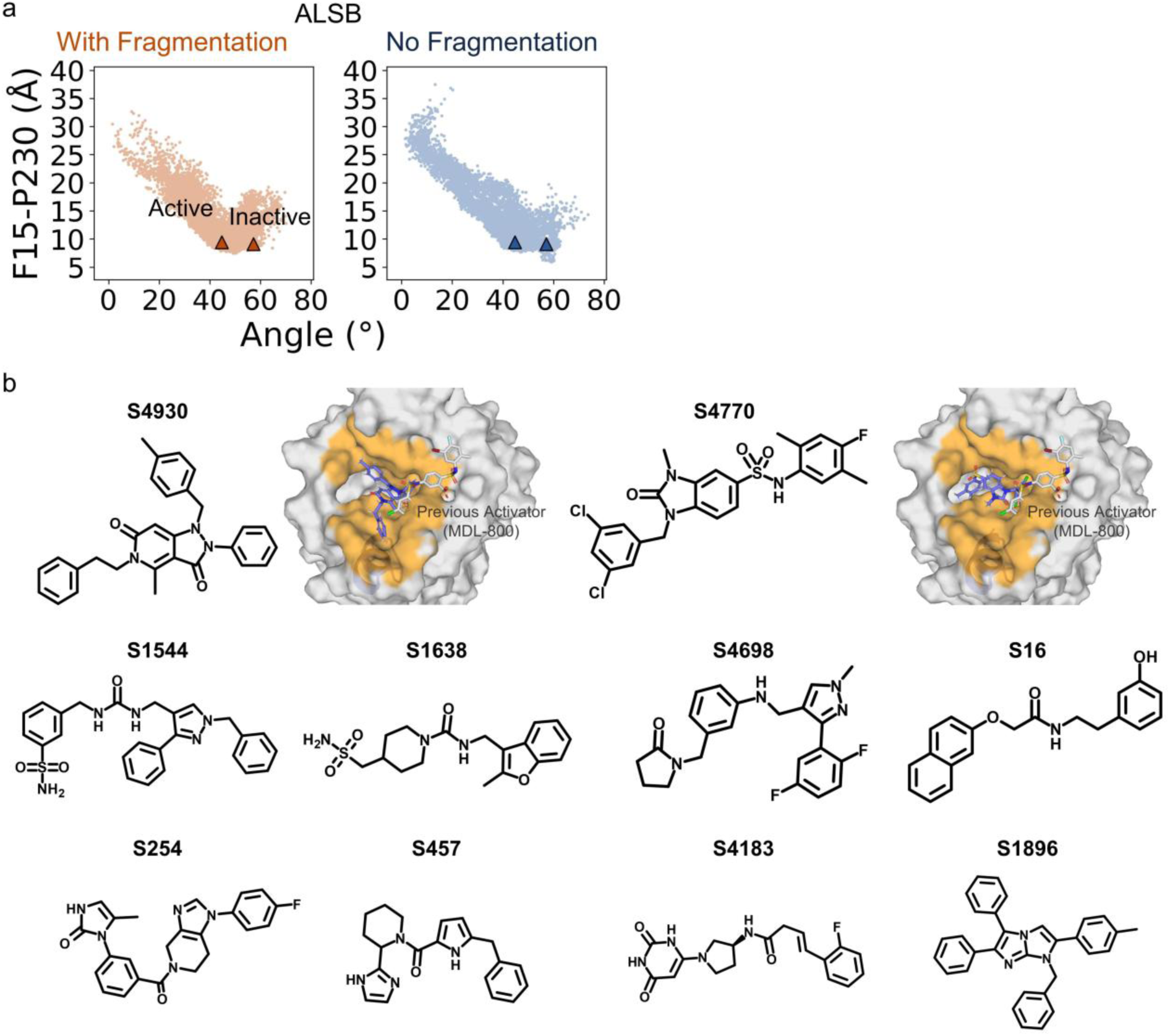
Sampling Active state conformations. **a.** Fragmentation doesn’t affect cases with only one domain moved. **b.** Top 10 selected compounds by virtual screening on the selected conformation of SIRT6. Docking poses of active compounds (blue) are given. A previous activator (white), MDL-800 is given for location reference.

**Extended Data Fig. 3.**
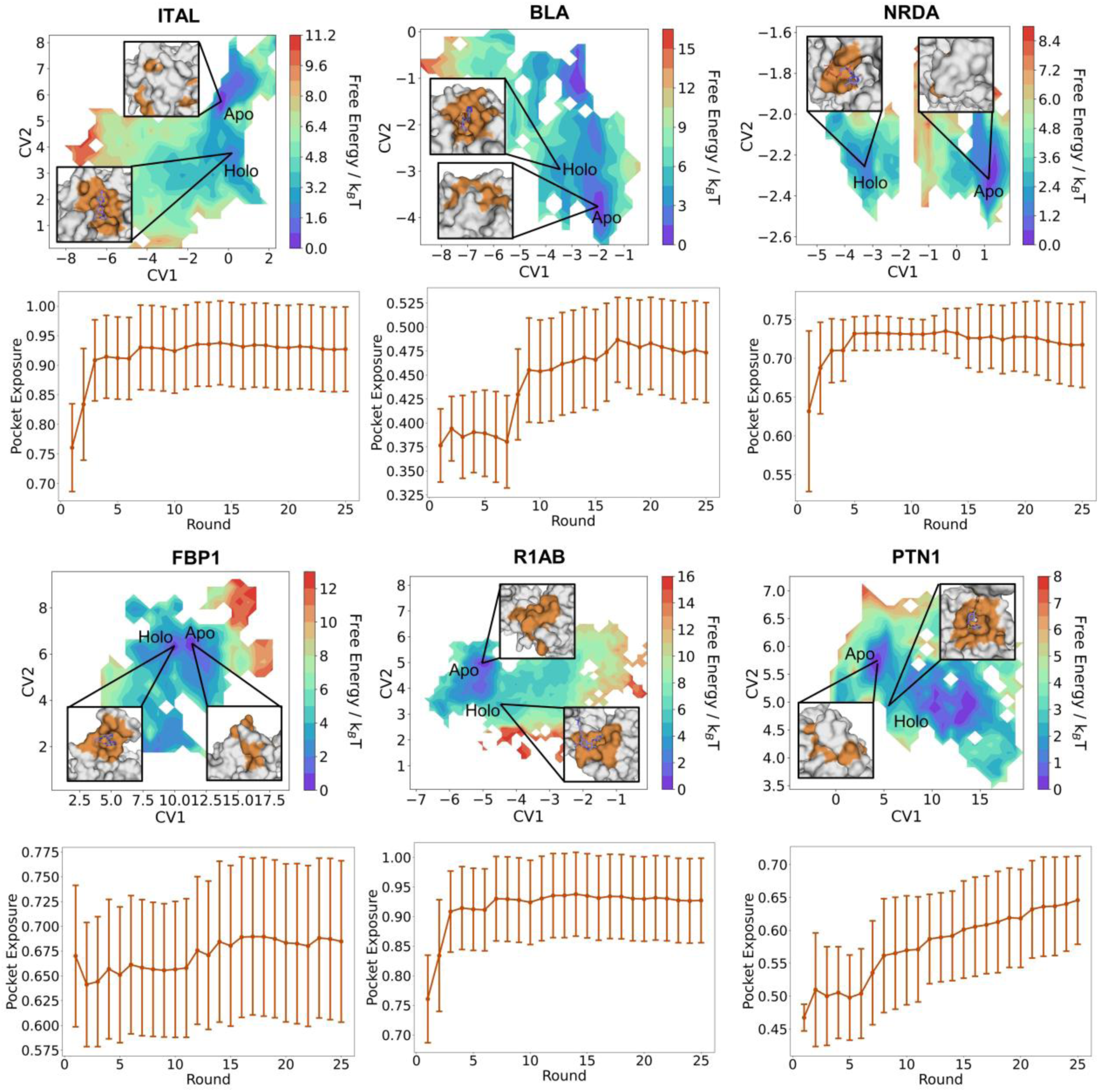
Sampling opening of cryptic allosteric sites. Top: Change of pocket exposure along iterations. Error bars represent standard deviations. Bottom: Dimension reduction by CTMAE with metastable states for open/closed conformations. N=400. Error bars represent standard deviations.

**Extended Data Fig. 4.**
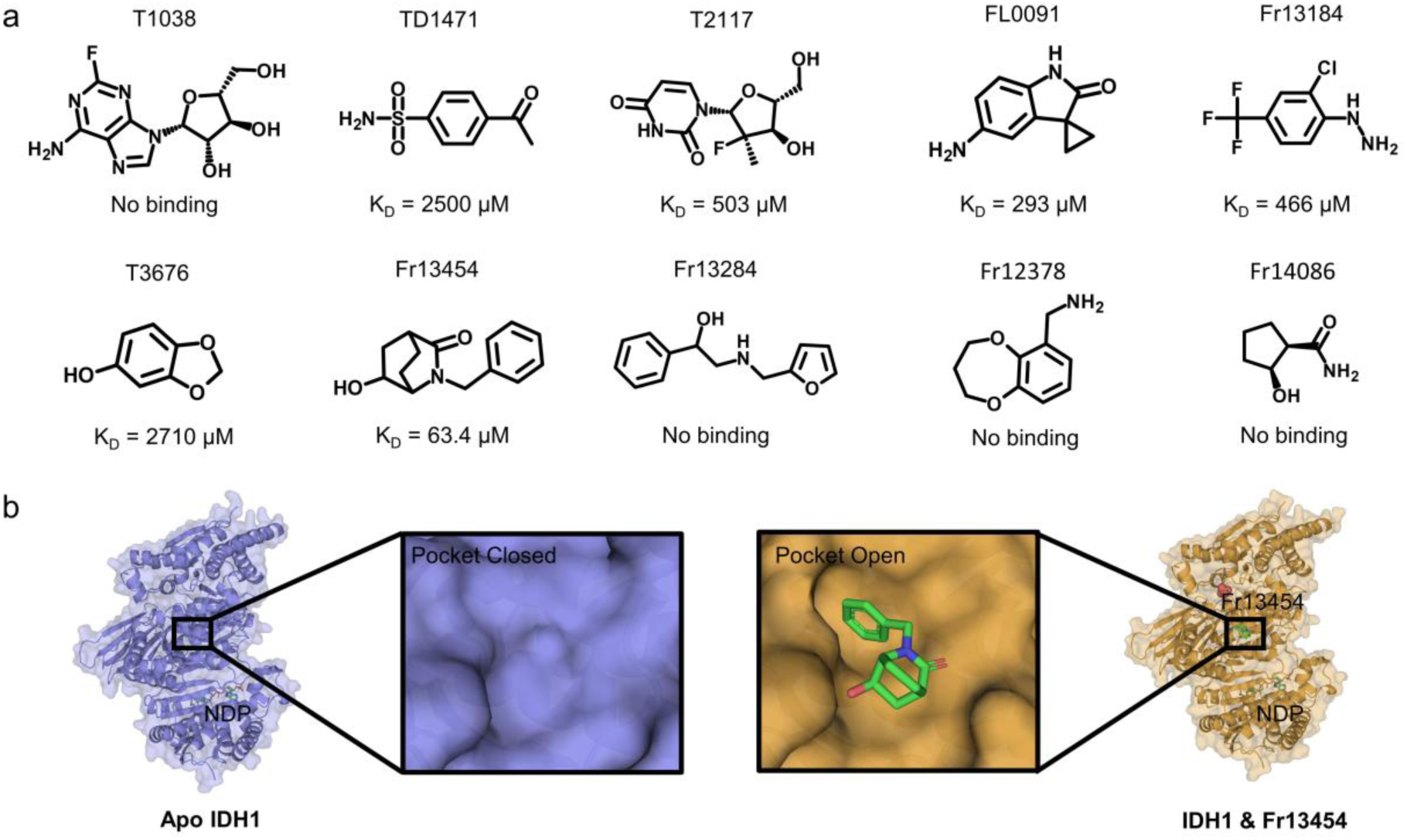
DASH Sampling of IDH1. **a.** Top 10 selected compounds by virtual screening on the selected allosteric pocket of IDH1. **b.** Comparison of the solved crystal structures of apo and Fr13454-bound IDH1. **c.** Distribution of helix degree calculated from 1000 samples generated by IDPSAM. Range of helix degree is set between 0 and the largest value (length of IDR minus 5).

**Extended Data Fig. 5.**
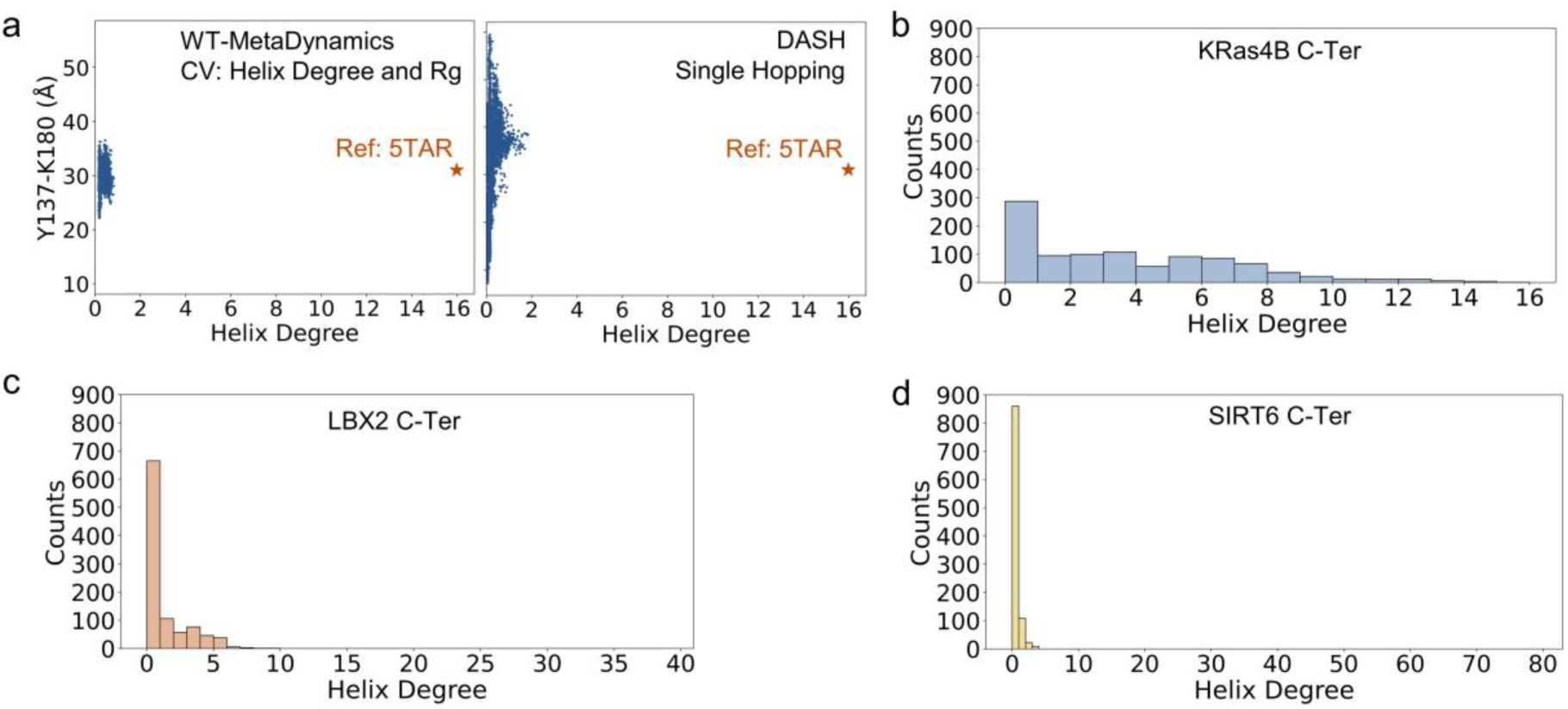
Sampling helical formation with combined methods. **a.** Sampling helix-folding of KRas4B HVR is failed using either well-tempered metadynamics (WT-Metadynamics) or DASH. **b-d.** Distribution of helix degree calculated from 1000 samples generated by IDPSAM. Range of helix degree is set between 0 and the largest value (length of IDR minus 5).

## Notes

### Competing Interest Statement

The authors have declared no competing interest.

## Reference

1. Ramelot, T.A., Tejero, R. & Montelione, G.T. Representing structures of the multiple conformational states of proteins. Curr. Opin. Struct. Biol. 83, 102703 (2023).

2. Sala, D., Engelberger, F., McHaourab, H.S. & Meiler, J. Modeling conformational states of proteins with AlphaFold. Curr. Opin. Struct. Biol. 81, 102645 (2023).

3. Saldaño, T. et al. Impact of protein conformational diversity on AlphaFold predictions. Bioinformatics 38, 2742–2748 (2022).

4. Olsson, U. & Wolf-Watz, M. Overlap between folding and functional energy landscapes for adenylate kinase conformational change. Nat. Commun. 1, 111 (2010).

5. Scheepstra, M. et al. Identification of an allosteric binding site for RORγt inhibition. Nat. Commun. 6, 8833 (2015).

6. Dharmaiah, S. et al. Structural basis of recognition of farnesylated and methylated KRAS4b by PDEδ. Proc. Natl. Acad. Sci. U.S.A. 113, E6766–E6775 (2016).

7. Rangari, V.A. et al. A cryptic pocket in CB1 drives peripheral and functional selectivity. Nature 640, 265–273 (2025).

8. De Vivo, M., Masetti, M., Bottegoni, G. & Cavalli, A. Role of Molecular Dynamics and Related Methods in Drug Discovery. J. Med. Chem. 59, 4035–4061 (2016).

9. Zhang, Q.F. et al. Targeting a cryptic allosteric site of SIRT6 with small-molecule inhibitors that inhibit the migration of pancreatic cancer cells. Acta Pharm. Sin. B 12, 876–889 (2022).

10. Ha, J.H. & Loh, S.N. Protein Conformational Switches: From Nature to Design. Chem. Eur. J. 18, 7984–7999 (2012).

11. Cavanagh, P.E. et al. Computational design of conformation-biasing mutations to alter protein functions. *Preprint at* https://www.biorxiv.org/content/10.1101/2025.05.03.652001v3 (2025).

12. Guo, A.B. et al. Deep learning-guided design of dynamic proteins. Science 388, eadr7094 (2025).

13. Klebl, D.P. et al. Swinging lever mechanism of myosin directly shown by time-resolved cryo-EM. Nature 642, 519–526 (2025).

14. Brändén, G. & Neutze, R. Advances and challenges in time-resolved macromolecular crystallography. Science 373, eaba0954 (2021).

15. Abramson, J. et al. Accurate structure prediction of biomolecular interactions with AlphaFold 3. Nature 630, 493–500 (2024).

16. Lin, Z.M. et al. Evolutionary-scale prediction of atomic-level protein structure with a language model. Science 379, 1123–1130 (2023).

17. Song, J., Ha, J., Lee, J., Ko, J. & Shin, W.H. Improving docking and virtual screening performance using AlphaFold2 multi-state modeling for kinases. Sci. Rep. 14, 25167 (2024).

18. Heo, L. & Feig, M. Multi-state modeling of G-protein coupled receptors at experimental accuracy. Proteins 90, 1873–1885 (2022).

19. Gkeka, P. et al. Machine Learning Force Fields and Coarse-Grained Variables in Molecular Dynamics: Application to Materials and Biological Systems. J. Chem. Theory Comput. 16, 4757–4775 (2020).

20. Mehdi, S., Smith, Z., Herron, L., Zou, Z.Y. & Tiwary, P. Enhanced Sampling with Machine Learning. Annu. Rev. Phys. Chem. 75, 347–370 (2024).

21. Kamenik, A.S., Linker, S.M. & Riniker, S. Enhanced sampling without borders: on global biasing functions and how to reweight them. Phys. Chem. Chem. Phys. 24, 1225–1236 (2022).

22. Hénin, J., Lelièvre, T., Shirts, M.R., Valsson, O. & Delemotte, L. Enhanced sampling methods for molecular dynamics simulations. *Preprint at* https://arxiv.org/abs/2202.04164 (2022).

23. Hu, F.C. et al. Unveiling the State Transition Mechanisms of Ras Proteins through Enhanced Sampling and QM/MM Simulations. J. Phys. Chem. B 128, 1418–1427 (2024).

24. Pan, Z.J., Li, M.D., Chen, D.C. & Yang, Y.I. A Sinking Approach to Explore Arbitrary Areas in Free Energy Landscapes. JACS Au 5, 2898–2908 (2025).

25. Fu, H.H., Shao, X.G., Chipot, C. & Cai, W.S. Extended Adaptive Biasing Force Algorithm. An On-the-Fly Implementation for Accurate Free-Energy Calculations. J. Chem. Theory Comput. 12, 3506–3513 (2016).

26. Li, M.Y. et al. Delineating the stepwise millisecond allosteric activation mechanism of the class C GPCR dimer mGlu5. Nat. Commun. 15, 7519 (2024).

27. Li, Y. & Gong, H.P. Identifying a Feasible Transition Pathway between Two Conformational States for a Protein. J. Chem. Theory Comput. 18, 4529–4543 (2022).

28. Lu, S.Y. et al. Activation pathway of a G protein-coupled receptor uncovers conformational intermediates as targets for allosteric drug design. Nat. Commun. 12, 4721 (2021).

29. Wang, J.N. et al. Gaussian accelerated molecular dynamics: Principles and applications. Wiley Interdiscip. Rev.: Comput. Mol. Sci. 11, e1521 (2021).

30. Wang, Y. et al. Delineating the activation mechanism and conformational landscape of a class B G protein-coupled receptor glucagon receptor. Comput. Struct. Biotechnol. J. 20, 628–639 (2022).

31. Xiong, Y.Q. et al. Conformations and binding pockets of HRas and its guanine nucleotide exchange factors complexes in the guanosine triphosphate exchange process. J. Comput. Chem. 43, 906–916 (2022).

32. Zeng, J. et al. Identification of functional substates of KRas during GTP hydrolysis with enhanced sampling simulations. Phys. Chem. Chem. Phys. 24, 7653–7665 (2022).

33. Kleiman, D.E., Nadeem, H. & Shukla, D. Adaptive Sampling Methods for Molecular Dynamics in the Era of Machine Learning. J. Phys. Chem. B 127, 10669–10681 (2023).

34. Tian, H. et al. LAST: Latent Space-Assisted Adaptive Sampling for Protein Trajectories. J. Chem. Inf. Model. 63, 67–75 (2023).

35. Chen, W. & Ferguson, A.L. Molecular enhanced sampling with autoencoders: On-the-fly collective variable discovery and accelerated free energy landscape exploration. J. Comput. Chem. 39, 2079–2102 (2018).

36. Sun, L. et al. Multitask Machine Learning of Collective Variables for Enhanced Sampling of Rare Events. J. Chem. Theory Comput. 18, 2341–2353 (2022).

37. Ribeiro, J.M.L., Bravo, P., Wang, Y. & Tiwary, P. Reweighted autoencoded variational Bayes for enhanced sampling (RAVE). J. Chem. Phys. 149, 072301 (2018).

38. Zhang, J., Yang, Y.I. & Noe, F. Targeted Adversarial Learning Optimized Sampling. J. Phys. Chem. Lett. 10, 5791–5797 (2019).

39. Trizio, E., Kang, P.L. & Parrinello, M. Everything everywhere all at once: a probability-based enhanced sampling approach to rare events. Nat. Comput. Sci. 5, 582–591 (2025).

40. Wang, D.D. et al. Efficient sampling of high-dimensional free energy landscapes using adaptive reinforced dynamics. Nat. Comput. Sci. 2, 20–29 (2022).

41. Zhang, M.Y., Wu, H. & Wang, Y. Enhanced Sampling of Biomolecular Slow Conformational Transitions Using Adaptive Sampling and Machine Learning. J. Chem. Theory Comput. 20, 8569–8582 (2024).

42. Shukla, D., Hernandez, C.X., Weber, J.K. & Pande, V.S. Markov state models provide insights into dynamic modulation of protein function. Acc. Chem. Res. 48, 414–422 (2015).

43. Dror, R.O. et al. Activation mechanism of theβ2-adrenergic receptor. Proc. Natl. Acad. Sci. U.S.A. 108, 18684–18689 (2011).

44. Scherer, M.K. et al. PyEMMA 2: A Software Package for Estimation, Validation, and Analysis of Markov Models. J. Chem. Theory Comput. 11, 5525–5542 (2015).

45. Ketkaew, R., Creazzo, F. & Luber, S. Machine Learning-Assisted Discovery of Hidden States in Expanded Free Energy Space. J. Phys. Chem. Lett. 13, 1797–1805 (2022).

46. Noe, F., Tkatchenko, A., Muller, K.R. & Clementi, C. Machine Learning for Molecular Simulation. Annu. Rev. Phys. Chem. 71, 361–390 (2020).

47. Chen, T., Kornblith, S., Norouzi, M. & Hinton, G. A Simple Framework for Contrastive Learning of Visual Representations. International Conference on Machine Learning (2020).

48. Palma, J. & Pierdominici-Sottile, G. On the Uses of PCA to Characterise Molecular Dynamics Simulations of Biological Macromolecules: Basics and Tips for an Effective Use. ChemPhysChem 24, e202200491 (2023).

49. Spiwok, V. & Králová, B. Metadynamics in the conformational space nonlinearly dimensionally reduced by Isomap. J. Chem. Phys. 135, 224504 (2011).

50. Pérez-Hernández, G., Paul, F., Giorgino, T., De Fabritiis, G. & Noé, F. Identification of slow molecular order parameters for Markov model construction. J. Chem. Phys. 139 (2013).

51. Dauparas, J. et al. Robust deep learning-based protein sequence design using ProteinMPNN. Science 378, 49–55 (2022).

52. Zhu, J.X., Wang, J.X., Han, W.W. & Xu, D. Neural relational inference to learn long-range allosteric interactions in proteins from molecular dynamics simulations. Nat. Commun. 13, 1661 (2022).

53. Gu, Z.H., Luo, X., Chen, J.X., Deng, M.H. & Lai, L.H. Hierarchical graph transformer with contrastive learning for protein function prediction. Bioinformatics 39, btad410 (2023).

54. Lu, S.Y. et al. Deactivation Pathway of Ras GTPase Underlies Conformational Substates as Targets for Drug Design. ACS Catal. 9, 7188–7196 (2019).

55. Do, P.C., Lee, E.H. & Le, L. Steered Molecular Dynamics Simulation in Rational Drug Design. J. Chem. Inf. Model. 58, 1473–1482 (2018).

56. Ramaswamy, V.K., Musson, S.C., Willcocks, C.G. & Degiacomi, M.T. Deep Learning Protein Conformational Space with Convolutions and Latent Interpolations. Phys. Rev. X 11, 011052 (2021).

57. Sun, Y.J., Rose, J., Wang, B.C. & Hsiao, C.D. The structure of glutamine-binding protein complexed with glutamine at 1.94 Å resolution: Comparisons with other amino acid binding proteins. J. Mol. Biol. 278, 219–229 (1998).

58. Wang, L., Friesner, R.A. & Berne, B.J. Replica Exchange with Solute Scaling: A More Efficient Version of Replica Exchange with Solute Tempering (REST2) (vol 115, pg 9431, 2011). J. Phys. Chem. B 115, 11305–11305 (2011).

59. Pasqualato, S., Ménétrey, J., Franco, M. & Cherfils, J. The structural GDP/GTP cycle of human Arf6. EMBO Rep. 2, 234–238 (2001).

60. Koshland, D.E. The Key-Lock Theory and the Induced Fit Theory. Angew. Chem. Int. Ed. 33, 2375–2378 (1994).

61. Xu, X.Y. et al. Discovery of a potent and highly selective inhibitor of SIRT6 against pancreatic cancer metastasis in vivo. Acta Pharm. Sin. B 14, 1302–1316 (2024).

62. Pan, P.W. et al. Structure and Biochemical Functions of SIRT6. J. Biol. Chem. 286, 14575–14587 (2011).

63. Huang, Z.M. et al. Identification of a cellularly active SIRT6 allosteric activator. Nat. Chem. Biol. 14, 1118–1126 (2018).

64. Chen, X.L. et al. Discovery of Potent Small-Molecule SIRT6 Activators: Structure-Activity Relationship and Anti-Pancreatic Ductal Adenocarcinoma Activity. J. Med. Chem. 63, 10474–10495 (2020).

65. You, W.J., et al. Structural Basis of Sirtuin 6 Activation by Synthetic Small Molecules. Angew. Chem. Int. Ed. 56, 1007–1011 (2017).

66. Lu, S.Y. et al. Mechanism of allosteric activation of SIRT6 revealed by the action of rationally designed activators. Acta Pharm. Sin. B 11, 1355–1361 (2021).

67. Zhao, Z.Y. et al. Deciphering the Allosteric Activation Mechanism of SIRT6 Using Molecular Dynamics Simulations. J. Chem. Inf. Model. 63, 5896–5902 (2023).

68. Ni, D. et al. Along the allostery stream: Recent advances in computational methods for allosteric drug discovery. Wiley Interdiscip. Rev.: Comput. Mol. Sci. 12, e1585 (2022).

69. Xie, J., Pan, G. & Lai, L. Sequence and Structure-based Prediction of Allosteric Sites. J. Mol. Biol., 169305 (2025).

70. He, J.X., et al. ASD2023: towards the integrating landscapes of allosteric knowledgebase. Nucleic Acids Res. 52, D376–D383 (2023).

71. Zhu, R.D., Wu, C.W., Zha, J.Y., Lu, S.Y. & Zhang, J. Decoding allosteric landscapes: computational methodologies for enzyme modulation and drug discovery. *RSC Chem*. Biol. 6, 539–554 (2025).

72. Lu, S.Y., Ji, M.F., Ni, D. & Zhang, J. Discovery of hidden allosteric sites as novel targets for allosteric drug design. Drug Discov. Today 23, 359–365 (2018).

73. Qiao, X. et al. Targeting cryptic allosteric sites of G protein-coupled receptors as a novel strategy for biased drug discovery. Pharmacol. Res. 212, 107574 (2025).

74. Ni, D. et al. Discovery of cryptic allosteric sites using reversed allosteric communication by a combined computational and experimental strategy. Chem. Sci. 12, 464–476 (2021).

75. Molenaar, R.J., Maciejewski, J.P., Wilmink, J.W. & van Noorden, C.J.F. Wild-type and mutated IDH1/2 enzymes and therapy responses. Oncogene 37, 1949–1960 (2018).

76. Khan, I., Waqas, M. & Shamim, M.S. Prognostic significance of IDH 1 mutation in patients with glioblastoma multiforme. J. Pak. Med. Assoc. 67, 816–817 (2017).

77. Le Guilloux, V., Schmidtke, P. & Tuffery, P. Fpocket: An open source platform for ligand pocket detection. BMC Bioinf. 10, 168 (2009).

78. Zha, J.Y. et al. AlloReverse: multiscale understanding among hierarchical allosteric regulations. Nucleic Acids Res. 51, W33–W38 (2023).

79. Hu, Y. et al. Exploring Protein Conformational Changes Using a Large-Scale Biophysical Sampling Augmented Deep Learning Strategy. Adv. Sci. 11, e2400884 (2024).

80. Pietrucci, F. & Laio, A. A Collective Variable for the Efficient Exploration of Protein Beta-Sheet Structures: Application to SH3 and GB1. J. Chem. Theory Comput. 5, 2197–2201 (2009).

81. Lu, S.Y., et al. Ras Conformational Ensembles, Allostery, and Signaling. Chem. Rev. 116, 6607–6665 (2016).

82. Ntai, I. et al. Precise characterization of KRAS4b proteoforms in human colorectal cells and tumors reveals mutation/modification cross-talk. Proc. Natl. Acad. Sci. U.S.A. 115, 4140–4145 (2018).

83. Parikh, K. et al. Drugging KRAS: current perspectives and state-of-art review. J. Hematol. Oncol. 15, 152 (2022).

84. Chakrabarti, M., Jang, H. & Nussinov, R. Comparison of the Conformations of Isoforms, K-Ras4A and K-Ras4B, Points to Similarities and Significant Differences. J. Phys. Chem. B 120, 667–679 (2016).

85. Zhang, H. et al. Markov State Models and Molecular Dynamics Simulations Reveal the Conformational Transition of the Intrinsically Disordered Hypervariable Region of K-Ras4B to the Ordered Conformation. J. Chem. Inf. Model. 62, 4222–4231 (2022).

86. Barducci, A., Bussi, G. & Parrinello, M. Well-tempered metadynamics: A smoothly converging and tunable free-energy method. Phys. Rev. Lett. 100, 020603 (2008).

87. Varadi, M. et al. AlphaFold Protein Structure Database: massively expanding the structural coverage of protein-sequence space with high-accuracy models. Nucleic Acids Res. 50, D439–D444 (2022).

88. Wang, J. et al. Identification of LBX2 as a novel causal gene of atrial septal defect *Int*. J. Cardiol. 265, 188–194 (2018).

89. Janson, G. & Feig, M. Transferable deep generative modeling of intrinsically disordered protein conformations. PLOS Comput. Biol. 20 (2024).

90. Qiu, Y.Y., Papai, G., Ben Shem, A. & Bignon, E. Specific Binding Modes of the SIRT6 C-Terminal Domain to the Nucleosome Core Particle Influence DNA Unwrapping and H3K27 Accessibility. J. Phys. Chem. B 129, 9855–9861 (2025).

91. Luo, P.Z. & Baldwin, R.L. Mechanism of helix induction by trifluoroethanol: A framework for extrapolating the helix-forming properties of peptides from trifluoroethanol/water mixtures back to water. Biochemistry 36, 8413–8421 (1997).

92. Culik, R.M., Abaskharon, R.M., Pazos, I.M. & Gai, F. Experimental Validation of the Role of Trifluoroethanol as a Nanocrowder. J. Phys. Chem. B 118, 11455–11461 (2014).

93. Kaczka, P. et al. The TFE-induced transient native-like structure of the intrinsically disordered domain of σ RNA polymerase. Eur. Biophys. J, 43, 581–594 (2014).

94. Ji, T. et al. Remote on-off switching of protein activity by intrinsically disordered region. Nat. Struct. Mol. Bio. 32, 2088–2098 (2025).

95. Liu, Y.L., Zhang, M.Z., Tsai, C.J., Jang, H. & Nussinov, R. Allosteric regulation of autoinhibition and activation of c-Abl. Comput. Struct. Biotechnol. J. 20, 4257–4270 (2022).

96. Liu, Y.L., Zhang, M.Z., Jang, H. & Nussinov, R. The allosteric mechanism of mTOR activation can inform bitopic inhibitor optimization. Chem. Sci. 15, 1003–1017 (2024).

97. Liu, Y.L., Zhang, W.G., Jang, H. & Nussinov, R. mTOR Variants Activation Discovers PI3K-like Cryptic Pocket, Expanding Allosteric, Mutant-Selective Inhibitor Designs. J. Chem. Inf. Model. 65, 966–980 (2025).

98. Case, D.A., et al. AMBER 2020. University of California, San Francisco (2020).

99. Eastman, P. et al. OpenMM 7: Rapid development of high performance algorithms for molecular dynamics. PLOS Comput. Biol. 13, e1005659 (2017).

100. Maier, J.A. et al. ff14SB: Improving the Accuracy of Protein Side Chain and Backbone Parameters from ff99SB. J. Chem. Theory Comput. 11, 3696–3713 (2015).

101. Jakalian, A., Jack, D.B. & Bayly, C.I. Fast, efficient generation of high-quality atomic charges. AM1-BCC model: II. Parameterization and validation. J. Comput. Chem. 23, 1623–1641 (2002).

102. Jakalian, A., Bush, B.L., Jack, D.B. & Bayly, C.I. Fast, efficient generation of high-quality atomic Charges. AM1-BCC model: I. Method. J. Comput. Chem. 21, 132–146 (2000).

103. Peters, M.B. et al. Structural Survey of Zinc-Containing Proteins and Development of the Zinc AMBER Force Field (ZAFF). J. Chem. Theory Comput. 6, 2935–2947 (2010).

104. Huang, J. et al. CHARMM36m: an improved force field for folded and intrinsically disordered proteins. Nat. Meth. 14, 71–73 (2017).

105. Lee, J. et al. CHARMM-GUI Input Generator for NAMD, GROMACS, AMBER, OpenMM, and CHARMM/OpenMM Simulations Using the CHARMM36 Additive Force Field. J. Chem. Theory Comput. 12, 405–413 (2016).

106. Eastman, P. et al. OpenMM 8: Molecular Dynamics Simulation with Machine Learning Potentials. J. Phys. Chem. B 128, 109–116 (2023).

107. de Souza, V.C., Goliatt, L. & Goliatt, P.V.Z.C. Clustering Algorithms Applied on Analysis of Protein Molecular Dynamics. Ieee Latin American Conference on Computational Intelligence (2017).

108. Paszke, A. et al. PyTorch: An Imperative Style, High-Performance Deep Learning Library. Adv. Neur. Inf. Process. Sys. (2019).

109. Ravindranath, P.A. & Sanner, M.F. AutoSite: an automated approach for pseudo-ligands prediction-from ligand-binding sites identification to predicting key ligand atoms. Bioinformatics 32, 3142–3149 (2016).

110. Lindorff-Larsen, K., Piana, S., Dror, R.O. & Shaw, D.E. How Fast-Folding Proteins Fold. Science 334, 517–520 (2011).

111. Xiao, Z.H. et al. A three-level regulatory mechanism of the aldo-keto reductase subfamily AKR12D. Nat. Commun. 15, 2128 (2024).

112. Shan, Y., Arkhipov, A., Kim, E.T., Pan, A.C. & Shaw, D.E. Transitions to catalytically inactive conformations in EGFR kinase. Proc Natl Acad Sci U S A 110, 7270–7275 (2013).

113. Research, D.E.S. Molecular Dynamics Simulations Related to SARS-CoV-2. D. E. Shaw Research Technical Data (2020).

114. Li, N. et al. Exploring the distinct activation mechanisms of neuromedin B receptor through multiple replica molecular dynamics simulations and Markov state modeling. Acta Pharmacol. Sin., doi: 10.1038/s41401-41025-01603-w (2025).

115. Trozzi, F., Wang, X.L. & Tao, P. UMAP as a Dimensionality Reduction Tool for Molecular Dynamics Simulations of Biomacromolecules: A Comparison Study. J. Phys. Chem. B 125, 5022–5034 (2021).

116. Lemke, T. & Peter, C. EncoderMap: Dimensionality Reduction and Generation of Molecule Conformations. J. Chem. Theory Comput. 15, 1209–1215 (2019).

117. Sainburg, T., McInnes, L. & Gentne, T.Q. Parametric UMAP embeddings for representation and semi-supervised learning. *Preprint at* https://arxiv.org/abs/2009.12981, 2009.12981 (2009).

118. Pedregosa, F. et al. Scikit-learn: Machine Learning in Python. J. Mach. Learn. Res. 12, 2825–2830 (2011).

119. Halgren, T.A. et al. Glide: A new approach for rapid, accurate docking and scoring. 2. Enrichment factors in database screening. J. Med. Chem. 47, 1750–1759 (2004).

120. Friesner, R.A. et al. Glide: A new approach for rapid, accurate docking and scoring. 1. Method and assessment of docking accuracy. J. Med. Chem. 47, 1739–1749 (2004).

121. Xiao, Q.J. et al. Upgrade of crystallography beamline BL19U1 at the Shanghai Synchrotron Radiation Facility. J. Appl. Crystallogr. 57, 630–637 (2024).

122. Kabsch, W. XDS. Acta Crystallogr., Sect. D: Biol. Crystallogr. 66, 125–132 (2010).

123. Agirre, J. et al. The CCP4 suite: integrative software for macromolecular crystallography. Acta Crystallogr., Sect. D: Biol. Crystallogr. 79, 449–461 (2023).

124. Adams, P.D. et al. PHENIX: a comprehensive Python-based system for macromolecular structure solution. Acta Crystallogr., Sect. D: Biol. Crystallogr. 66, 213–221 (2010).

125. Afonine, P.V. et al. Towards automated crystallographic structure refinement with phenix.refine. Acta Crystallogr., Sect. D: Biol. Crystallogr. 68, 352–367 (2012).

126. Emsley, P., Lohkamp, B., Scott, W.G. & Cowtan, K. Features and development of. *Acta Crystallogr., Sect. D: Biol*. Crystallogr. 66, 486–501 (2010).

